# Phase-locking saccades to posterior alpha oscillations improves the neural representation of visual objects during memory formation

**DOI:** 10.1101/2025.10.09.681394

**Authors:** Graham Flick, Jed Meltzer, Jennifer D. Ryan, Rosanna K. Olsen

**Author notes:** Corresponding Author: Graham Flick.

## Abstract

Visual memory formation begins with the intake and neural processing of discrete samples provided by gaze fixations and saccades. Past research has highlighted a functional relationship between the timing of saccades and oscillations in neural activity over posterior brain areas: saccades phase-lock to alpha oscillations (8-12 Hz) and the degree of phase-locking, in natural scenes, has been associated with subsequent recognition memory. Here, we tested the hypothesis that the putative memory encoding benefit arises due to improved neuronal processing and, ultimately, better neural representation of foveated items, when saccades are locked to alpha phase. In a co-registered magnetoencephalography (MEG) and eye-tracking paradigm, participants executed saccades from central fixation to images that appeared in the periphery, attempting to remember those images for later testing. Replicating past results, saccades to subsequently remembered images were preceded by greater inter-trial phase coherence in the alpha frequency band, at posterior MEG channels, consistent with the notion that the eye movements were phase-locked. Across participants, the degree of saccade phase-locking was positively correlated with how well visual and semantic properties of the images were represented in neural activity, within 200 ms of their foveation. This relationship was evident in responses localized to the left parieto-occipital and ventral temporal cortex, where greater saccade phase-locking was associated with improved visual and semantic representations, respectively. These results support the hypothesis that phase-locking saccades to alpha oscillations leads to improved neuronal representation of foveated stimuli, providing new mechanistic insight into how episodic memories are formed from discrete visual samples.

**Significance Statement:** Visual memories begin with discrete samples provided by eye movements and gaze fixations, from which the brain must extract information that is integrated into a coherent memory trace. This work examined the neural mechanisms that support this memory formation process, one eye movement at a time. The results identified a functioning coupling between saccade timing and neuronal oscillations in the alpha frequency band: when saccades are phase-locked to alpha oscillations, the brain can better process the visual information provided by gaze fixations. This indicates that precise coordination of saccade timing, with respect to neuronal oscillations, promotes more veridical neural representation of the resulting visual samples, from which accurate episodic memories can be constructed.

## 1. INTRODUCTION

The formation of visual episodic memories relies on perceptual input provided by eye movements and gaze fixations, bound together to form coherent memory traces (Voss et al., 2017; Ryan, Shen, & Liu 2020). Neuroscientific evidence indicates that this process involves tight coordination between saccades – rapid, ballistic eye movements – and neuronal oscillations in the alpha (8-12 Hz) frequency range (Staudigl et al., 2017; Liu, Nobre, & van Ede, 2022; Popov & Staudigl, 2023). Here, we provide new evidence that saccade timing, relative to alpha oscillations, impacts memory formation by influencing how well fixated visual objects are represented in brain activity.

Dating back to Berger (1929), there has been an association between the power of alpha oscillations and visual processing, predominantly examined in passive viewing paradigms (i.e., presentation of stimuli during fixation; Ergenoglu et al., 2004; Hanslmayr et al., 2005; van Dijk et al., 2008). Beyond power, alpha phase has also been shown to modulate visual cognition, influencing the detection of briefly presented targets (Busch, Dubois, & VanRullen, 2009; Matthewson et al., 2009) and the relay of cortical responses to visual stimuli or phosphene-inducing neural stimulation (Dugué, Macque & VanRullen, 2011; Hanslmayr et al., 2013). These findings align with accounts of alpha oscillations as pulses of inhibition (Jensen & Mazaheri, 2010; Matthewson et al., 2011), such that stimuli presented at optimal phases receive improved neuronal processing.

More recent findings highlight a role of alpha oscillations in active vision as well. When viewers explore scenes, the brain’s visual and dorsal attention networks exhibit prominent alpha-band oscillations, which support interaction with the hippocampus during memory encoding (Kragel et al., 2021). Several studies have also found that saccades phase-lock to alpha oscillations (Drewes & VanRullen, 2011; Staudigl et al., 2017; Pan et al., 2023). Most notably, Staudigl and colleagues (2017) reported that memory for scenes was related to the degree of saccade phase-locking: when saccades were executed at consistent alpha phases, the encoded scenes were more likely to be recognized in a subsequent memory test. This suggests that eye movements were timed to promote memory formation, but greater insight is needed into how this benefit arises, mechanistically.

Based on the findings that alpha phase influences stimulus detection (Matthewson et al., 2009), as well as past work demonstrating the role of alpha phase in inter-areal neural communication (Dugue et al., 2011; Kragel et al., 2021; and see Voloh & Womelsdorf, 2016), we hypothesized that phase-locking saccades to alpha improves the neuronal processing of foveated visual elements within a scene, resulting in an improved neural representation of those elements, from which the memory can be constructed. From this, we predicted that when participants show heightened saccade-alpha phase-locking, their neural representations of foveated stimuli would display greater representational fidelity, captured in multivariate pattern analyses.

We tested these predictions in a co-registered MEG and eye-tracking study. During an initial encoding phase, participants made saccades to visual items that appeared in the periphery, attempting to remember them for later testing (Figure 1). We then back-sorted trials as subsequent hits and misses and analyzed neural activity time-locked to saccades to the encoded items. While past studies have characterized neural representations in brain activity after image presentation (e.g., Carlson et al., 2011, 2013; Clarke et al., 2018), very little research has examined how these representations emerge after eye movements (e.g., Fakche et al., 2024).

**Figure 1.**
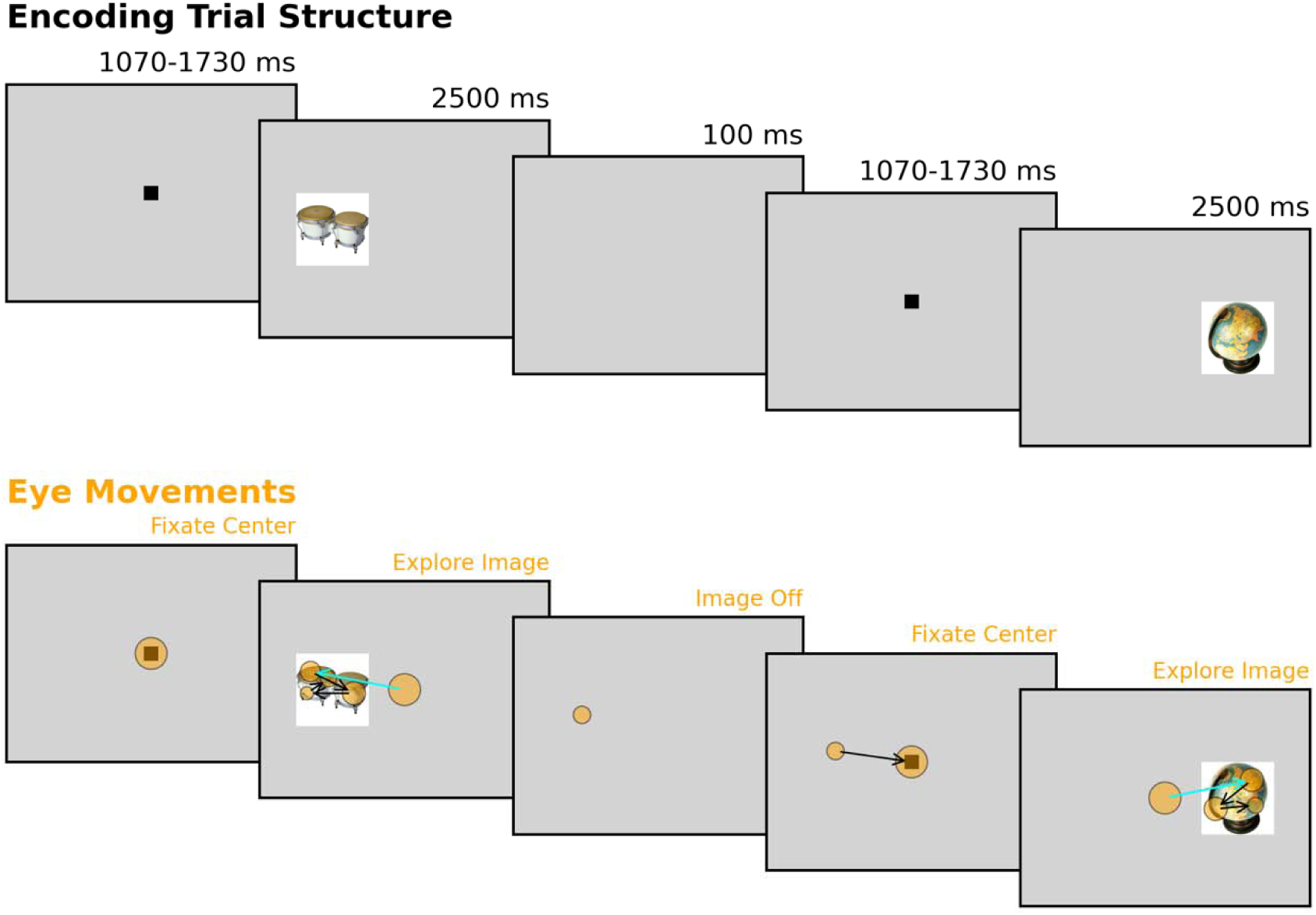
Structure of the encoding task, showing duration of each event (top) and example eye-tracking data (bottom). Each orange circle indicates a gaze fixation, with the radius scaled by duration. Arrows indicate saccades. The cyan arrow indicates the first saccade to the image, after it appeared on screen. Images are not to scale.

Our results identified memory-related differences in saccade phase-locking in the alpha frequency band, matching previous results, and demonstrated that greater phase-locking was associated with improved neural representation of the visual and semantic properties of newly fixated objects. This relationship was unique to pre-saccade phase (i.e., not power) and was present in activity localized to left parieto-occipital cortex and the ventral temporal lobe. These results support the hypothesis that phase-locking saccades to alpha oscillations leads to improved neuronal representation of foveated stimuli and provide new mechanistic insight into how this coupling facilitates memory formation.

## 2. MATERIALS AND METHODS

### 2.1 Participants

Thirty-six participants between the ages of 18 and 40 years were recruited from the Greater Toronto Area. This sample size was chosen to match that of the previous research directly motivating this investigation (Staudigl et al., 2017). All participants were fluent English speakers, with normal hearing and normal vision, or corrected-to-normal vision with the use of soft contact lenses. Exclusion criteria included a history of neurological, mood, anxiety, psychological, or substance abuse disorders assessed via self-report, and any non-removable metal in or on the body. Four participants who were recruited did not complete the experimental procedures due to issues at MEG data collection, including the identification of non-removable metal and malfunctions of the MEG acquisition system. The final sample of 32 participants included 23 females and 9 males, with a mean age of 23.4 years (standard deviation = 4.40 years, range = 19-37 years).

### 2.2 Experimental Design and Stimuli

The study consisted of three primary phases: (i) a memory encoding task; (ii) a distractor task consisting of a 9-minute short-film; and (iii) a memory recognition task. These were preceded and followed by two additional tasks: a saccade task prior to the initial encoding phase (5 minutes), and a memory assessment for the film (5 minutes). In the current paper, we focus exclusively on MEG and eye-tracking data collected during the memory encoding.

In the encoding task, participants were sequentially presented with 250 unique images and asked to remember as many of them as possible for the subsequent recognition memory assessment. Each image was presented once at encoding, on either the left or right side of the screen, such that participants were required to make a saccade from central fixation to view the image. In the recognition task, participants saw the original 250 images interspersed with an additional 250 novel images. All images were presented centrally during the recognition task.

Images were extracted from the DinoLab database (Hovhannisyan et al., 2021) using 22 of the available semantic categories (furniture, musical instruments, plants, street items, household items, food, vehicles, electronics, tools, body parts, insects, mammals, fruits, buildings, clothing, vegetables, sports items, kitchen items, toys, sea objects, birds, and decorative items). Half of the images in each semantic category were assigned to the Target set, presented at encoding, whereas the other half were assigned to the Foil set, presented only at recognition. This was done so that the image category could not be a reliable cue in the recognition memory test. Within each semantic category, images were selected and assigned to Targets and Foils to minimize the difference between the average visual hit and false alarm rates reported in the memory norming experiments by Hovhannisyan et al. (i.e., the successful recognition of the image when tested a day later). This was done to reduce potential response biases at recognition testing. The median hit rates extracted from the norming data of the Targets and Foils were 57.4% and 58.7%, respectively, while the median false alarm rates were 25.9% and 23.8% respectively, indicating minimal differences.

During the distractor phase, participants watched the short film *Growth* (https://www.silvanderwoerd.com/growth) while seated in the MEG system. This approximately 9-minute film follows the life of a family of four in their home as the characters grow older, in a style that mimics a continuously panning camera. All spoken language in the film is in Dutch, which none of the participants reported being able to speak or understand. No subtitles were shown during the film.

### 2.3 Procedure

All participants provided informed consent before taking part in the study procedures. All procedures were approved by the Research Ethics Board of Baycrest Hospital and took place in the MEG and magnetic resonance imaging (MRI) laboratories of Baycrest Hospital.

Before they entered the magnetically shielded room (MSR) that housed the MEG system, each participant was provided an overview of the three phases of the study, using example images that were not included in the subsequent procedures. Participants were shown example trials from both the encoding and recognition phases, and were informed that their memory of the images encountered during the encoding phase would be tested in later visual recognition trials. They were told half of the images at test would be new and the other half would be the original set, shown at encoding. Experimenters then recorded each participant’s fiducial landmarks (nasion and left/right tragi), as well as the position of three head position indicator coils placed on the participant, using digital photographs.

Participants sat upright in the MEG system during the experiment, with an eye-tracker placed at chest-level on a plastic mount in front of them, approximately 70 cm away. A visual back-projection screen was placed behind and above the eye-tracker, so that no part of the projected image was occluded. The screen was positioned approximately 85 cm from the participant’s eyes, with individual distances varying approximately 5-10 cm depending on the position of the participant in the MEG system. The projected display resolution was 1920 x 1080 pixels and the projected image spanned approximately 45 x 26 cm, in the horizontal and vertical directions, respectively. Prior to beginning the experiment, the experimenter confirmed that the eye-tracker maintained tracking of the pupil and corneal reflection while the participant looked at all four corners of the screen.

The experiment began with a five-point horizontal and vertical eye-tracking calibration, performed using the default HV5 EyeLink routine. Calibrations were accepted if the average point error was less than 1 degree, with no single point showing a margin of error worse than 1.5 degrees. The calibration-validation routine was repeated between every run of the encoding and retrieval phases of the experiment. If calibration failed multiple times, participants were given a short (10-15 second) break to rest their eyes. The next run began only after the eye-tracking calibration met the accuracy threshold.

The encoding phase of the study was divided into five runs of 50 trials. Each trial began with a centrally presented fixation square shown for 1400 milliseconds (ms) plus a random uniformly sampled jitter ranging from plus/minus 330 ms, totaling 1070 to 1730 ms. Participants were instructed to fixate on the square at the start of every trial. At the same time as the disappearance of the square, a 400 x 400 pixel image, subtending 6.25 degrees of visual angle (dva), appeared on either the left or right side of the screen. The center of each image was offset approximately 7.45 dva from the center of the screen.

Images were pseudo-randomly assigned to the left or right side of the screen for each participant, such that no more than 4 consecutive trials displayed an image on the same side. Each image was shown on screen for 2500 ms before removal. Participants were instructed to freely view each image and attempt to remember it for later testing. On 90% of trials, this was followed by a 100 ms blank screen before the central fixation square reappeared to mark the start of the next trial. On the remaining 10% of trials, participants were asked a probe question about the object that was just on the screen, which served as an attention check and encouraged semantic processing of the depicted items. This probe appeared in the center of the screen 100 ms after the offset of the to-be-encoded image, and asked one of three questions: (i) Was the item ever alive? (ii) Is the item typically found indoors or outdoors? and (iii) Does the item fit in a shoebox?

Participants responded “Yes” or “No” to each probe using a Current Designs (Philadelphia, PA) MEG-compatible response box with their right hand. Each probe question remained on the screen until a response was made. An equal number of each probe question was randomly assigned to items for each participant and trials on which the probes occurred were excluded from MEG analyses related to memory performance for that participant. Each run of the encoding task took approximately 5 minutes for participants to complete, with the total encoding task time, including eye-tracking calibration and validation before each run, requiring approximately 30 minutes.

Following completion of the encoding phase, participants viewed the short film in the distractor phase of the study. Participants were instructed to pay close attention to the film. The film’s presentation was preceded by calibration and validation routines of the eye-tracking system. This resulted in approximately 10-15 minutes of delay between the end of the encoding phase and the start of the memory recognition phase.

The recognition task was divided into 5 runs, each consisting of 100 trials. The order of the 500 images was randomly determined for each participant. Each trial began with a central fixation square that was displayed on screen for 1000 ms plus a jitter randomly sampled from a uniform distribution of plus/minus 83 milliseconds, for a possible total duration of 917-1083 ms. A 400 x 400 pixel image (subtending 6.25 dva) was then presented at the center of the screen for 2000 ms. Upon its removal, the next screen prompted participants to judge whether the image was previously encountered in the encoding phase of the experiment (an “Old” response) or not (a “New” response). Subsequently, they were asked to rate their confidence in their judgment from 1 to 4, where 1 indicated that they were just guessing and 4 indicated that they were certain they encountered the image at encoding. Each response display remained on screen until the participant pressed a button to indicate their decision. Each run of the recognition phase required approximately 10 minutes for participants to complete.

All experimental tasks were programmed in Psychopy (v. 2023.1; Peirce et al., 2019) in the Python computing environment (v. 3.11.5) and run on a Dell Latitude E6440 laptop. The EyeLink system was controlled from this laptop via ethernet cable connection using the application programming interface provided by SR Research in the EyeLink Developers kit and the PyLink Python wrapper. Visual stimuli were back projected to the participant’s viewing screen via a series of two mirrors using a PROPixx visual projector (VPixx, Saint-Bruno, QC) mounted outside of the magnetically shielded room.

### 2.4 Data Collection

MEG data were collected using a 275-channel, whole head, axial gradiometer CTF system (CTF, Coquitlam, British Columbia) with an online low-pass filter of 600 Hz and a sampling rate of 1200 Hz. Before and after each run of the MEG experiment, the participant’s head position was localized by sending electrical current to three head position indicator coils placed on the participant’s nasion and beside each tragus. A small towel was placed in the MEG helmet to provide a snug fit around the participant’s head in order to minimize movement, while keeping the scalp as close to the MEG sensors as possible. Eye-tracking data were collected using an EyeLink 1000 MEG-compatible long-range system, with monocular tracking of the right eye and a sampling rate of 1000 Hz. For all but two participants, a high-resolution (192 × 256 × 160 voxels; 1 mm^3^ isotropic voxels) T1-weighted MP-RAGE structural magnetic resonance imaging (MRI) scan was collected from each participant on a 3-Tesla system (Siemens Magnetom Prisma) located at Baycrest Hospital. For the remaining two individuals, a previously collected 3-Tesla T1 scan, with similar voxel size and image resolution (224 x 320 x 320; 0.8 mm^3^ voxels), was used in source localization.

The timing of each central fixation square and image onset during the encoding and recognition phases of the experiment was recorded in the continuous MEG data and used to subsequently define epochs of data for analysis. This utilized the Propixx projector’s transistor-transistor logic (TTL) digital output capability to send TTL pulses to the MEG system, which were recorded on an auxiliary channel alongside the MEG data. These TTL pulses, or triggers, encoded the colour of the top left-most pixel in the projected image on a frame-by-frame basis. The colour was programmed to change simultaneously with any event of interest in the experiment, thereby defining the onset (and offset) of that event in the MEG recording.

Two methods were used to ensure accurate and precise co-registration of the MEG and eye-tracking data recordings. First, the experiment control program sent a message indicating the onset of every event of interest (fixation squares, images, response screens) to the EyeLink Host computer, simultaneously with the display of the image. These messages were sent via the control computer’s ethernet connection with the EyeLink host computer and were recorded alongside the continuous eye position in the .EDF EyeLink output file. Second, an analog output card in the EyeLink system sent the live sample-by-sample position of the tracked eye, in x- and y-gaze coordinates, to the MEG system, where it was recorded on auxiliary channels alongside the continuous MEG data.

The onset times of all saccades, fixations, and blinks were initially defined from the .EDF output file using the standard EyeLink parser. We then defined the corresponding time point of each eye movement event in the MEG data using the relative time, after adjusting for the difference in sampling rates, from the onset of the fixation square or image on that trial, which was recorded in both data modalities. To adjust for any misalignments, we extracted the continuous samples of the x- and y-gaze position data from the .EDF EyeLink file and from the auxiliary MEG channels. Using the saccade and fixation onset times from the .EDF file relative to the start of each trial, we extracted 125 samples before/after each event in the eye-tracking data, as well 150 samples before after/each corresponding event in the MEG data. We then computed the cross-correlation between the two signals and extracted the Pearson correlation coefficients for each time lag relative to the onset time of the eye movement event defined in the MEG data from the .EDF file. This was done separately for both the x- and y-gaze position time series and the onset time of the eye movement event was then redefined in the MEG data at the time point with the maximum correlation coefficient in the local time window. This identifies the precise start and end time of eye movement events in the MEG data, using the event detection tools applied to the .EDF file.

### 2.5 Statistical Analysis

#### 2.5.1 MEG Data Preprocessing

The continuous MEG data were high- and low-pass filtered at 1 Hz and 40 Hz respectively. The data were then visually inspected and segments containing high levels of electromagnetic noise were annotated for removal. In two participants, the final two runs of the encoding task were lost due to a malfunction of the MEG acquisition system, resulting in these participants providing only three runs of MEG encoding data. Independent component analysis (ICA) was used to remove artifacts related to eye-blinks, saccades, cardiac activity and muscle contractions. Saccade-related components were further inspected by applying the same ICA solution to each participant’s saccade-related epochs (see definition below) to confirm that the component’s salience was highest at the onset of eye movements. Subsequently, a second ICA was applied to decompose each participant’s epochs of saccade-locked data (-1000 to 1000 ms surrounding all saccade onsets) to identify and remove any remaining components that reflected saccadic artifacts.

The cleaned data were segmented into two sets of epochs: image-related epochs spanning +/- 1000 ms around the onset of image and saccade-related epochs spanning +/- 1000 ms around the onset of each saccade. Each epoch was annotated with information about the side of the screen the image appeared on, whether the image was correctly recognized at subsequent test and, in the case of saccade-locked epochs, several properties related to the eye movement: the onset and offset latencies, duration, amplitude, and gaze position of the eye at launch and landing. We focused our primary analyses on the first saccade to the images during encoding. This was defined as the saccade with the earliest onset time following the appearance of the image, which landed on the image while it was on screen, was not preceded by any earlier fixations on the image after its appearance, and was launched from a 300 x 300 pixel rectangle in the center of the screen. The last two criteria eliminated any trials on which participants were already foveating the region in which the image appeared, instead of maintaining central fixation.

#### 2.5.2 Time-frequency Analysis: Inter-trial phase coherence

Time-resolved power and phase were estimated from image- and saccade-related epochs using complex Morlet wavelet convolution at 2 Hz frequency bins from 4 to 30 Hz, with two cycles defining the width of each wavelet’s Gaussian taper. Inter-trial phase coherence (ITPC; Tallon-Baudry et al., 1996) was used to assess the consistency of phase-angles preceding the execution of the saccades, where ITPC is derived from the circular sum of the phase-angles across trials calculated at each discrete time point, and ranges from 0 to 1, with 1 indicating perfect phase coherence.

Using the saccade-locked epochs, we calculated each participant’s time-course of mean ITPC values from the set of 44 bilateral parietal MEG channels (MLP/MRP in the CTF-275 system), then averaged those across the alpha frequency band (8-12 Hz). In the primary analyses, ITPC was baseline corrected using a 100 ms interval extracted from 500 ms before saccade onset, during the central fixation window, however the alpha-band ITPC effects (both the contrast of hits and misses, and the relationship with the RSA coefficients) were robust across multiple choices of baseline correction windows, including 100 ms intervals beginning 400 and 300 ms before saccade onset, and when no baseline correction was applied. To test whether we could replicate the previous finding of greater saccade-alpha phase-locking related to better subsequent memory (Staudigl et al., 2017), we compared the ITPC values calculated from the epochs that were later remembered to those that were later forgotten. To ensure that differences in trial numbers did not bias the estimate of ITPC, we first equated the number of trials in each condition. Epochs were dropped from the condition (hits or misses) that had a greater number of trials. The dropped trials were selected in a manner that made the remaining sets of epochs as close in time as possible, within the overall recording session (using the mne-python function epochs.equalize_event_counts, with method equal to “mintime”). Difference waves were then calculated by subtracting each participant’s alpha-band ITPC for subsequent misses from that for subsequent hits.

Group-level analysis of the ITPC was performed using a cluster-based permutation test (Maris & Oostenveld, 2007) on the 300 ms time courses of alpha-band ITPC differences preceding the onset of saccades, chosen to capture the time window of previous effects (Staudigl et al., 2017). Difference values were converted to a time course of t-statistics before temporal clusters were formed from any contiguous time points that corresponded to a p-value of less than 0.05. Cluster-level statistics were calculated as the sum of the encompassing t-values and compared to a surrogate null distribution generated from 5000 random sign-flips of the ITPC difference waves. If any true cluster-level statistic was greater than 95% of the surrogate null distribution, it was considered statistically significant evidence against the null hypothesis of no difference. Following the recommendations of van Diepen & Mazaheri (2018) and Nikolaev et al. (2013, 2016), we performed several follow-up analyses to ensure that pre-saccade differences in ITPC were not related to underlying differences in signal-to-noise ratios when estimating phase angles, or due to artifacts stemming from short previous fixations.

#### 2.5.3 Representational Similarity Analysis

Representational similarity analysis (RSA; Kriegeskorte, Mur, & Bandettini, 2008) was used to quantify the emergence of visual and semantic representations in distributed patterns of neural activity, recorded outside of the head by the MEG sensors and subsequently localized to each participant’s cortical surface. RSA compares the pairwise (dis)similarity between stimuli derived from a model, with that which emerges in pairwise comparisons of neural activity in response to those stimuli. If the two (dis)similarity spaces are themselves similar to one another, beyond that expected by chance alone, it supports the inference that the examined neural activity encodes those representations.

Here, we defined three relevant levels of representation. The first two, referred to as early and late visual representations, were defined using the AlexNet convolutional neural network model for image classification (Krizhevsky, Sutskever, & Hinton, 2017). Previous work using the same early and late visual definitions has demonstrated that vectorized representations of images extracted from the fully-connected layers of the AlexNet model correspond with representations found in human brain activity during visual processing (Clarke, Devereux, & Tyler, 2018; Devereux, Clarke, & Tyler, 2018). We used the *things-vision* Python package (Muttenthaler & Hebart, 2021) to extract the vectors of activation in the 3rd-to-last (early visual representation) and 2nd-to-last (late visual representation) fully connected layers of the model, when each image was passed through its pre-trained weights. We then computed model-based dis-similarities from each pair of images by subtracting the Pearson correlation coefficient of the two vectors from 1. When arranged in matrix-form, this provided representational dis-similarity matrices (RDMs) for the early and late visual representations.

We also computed a semantic-level model dis-similarity matrix using semantic feature production norming data. These data were available alongside the image dataset from which we selected our stimulus set (Hovhannisyan et al., 2021). Beginning with the set of 995 publicly available images and their feature vectors, we first set any feature that was produced by less than 3 participants for a given image to zero, in order to reduce noise in the feature vectors. We then used principal component analysis (PCA) to truncate the original 5020-dimensional feature space to 30 dimensions. This choice of dimensionality was motivated by inspection of the total explained variance with each additional dimension, which showed a drop in marginal explained variance at approximately 30 components. Using the truncated vectors, we computed a model-based RDM for semantic representations by calculating the pairwise Pearson correlation coefficient between each pair of vectors and subtracting those coefficients from 1.

Neural RDMs were computed for each participant at the MEG sensors by vectorizing the recorded magnetic flux values in 50 ms sliding time windows, centered on every individual time point beginning 25 ms into the epochs (see Carlson et al., 2011, 2013, Clarke et al., 2018 for similar approaches). For each pair of images, the dis-similarity between a participant’s vectorized MEG responses was computed using 1 minus the Pearson correlation coefficient. This was repeated for every pair of presented images, at every time point in the MEG response, resulting in time courses of neural dis-similarity matrices, wherein each element in each matrix represents the dis-similarity between a pair of images at that time point. For each neural RDM, we then extracted and vectorized the upper triangle from the matrix and calculated the Spearman rank correlation coefficient with the vectorized upper triangle of each model RDM (i.e., early visual, late visual, semantic). This yielded a time-course of representational similarity values, for each participant, for each level of representation. We refer to these values, which we consider to quantify the representational fidelity of the viewed stimulus, as RSA coefficients.

The same RSA procedure was repeated in our analysis of source-localized MEG responses (see below, Section 2.6). There, instead of vectorizing the magnetic flux responses across the MEG channels, we vectorized the estimated activity across the constituent sources in each region of interest. The same routines were then used to derive pairwise dis-similarities in neural responses to each image, for each participant, and to perform group-level statistical analysis on source estimates.

Group-level analysis was then performed on the time courses of RSA coefficients using cluster-based permutation tests (using the same parameters specified in Section 2.5.2) in the 500 milliseconds following event onsets; a window selected to capture the emergence of visual- and semantic-based representations of visual stimuli in past studies (Carlson et al., 2011; 2013; Clarke et al., 2018), including those that have examined neural representations following eye movements (Auerbach-Asch & Deouell, 2024; Fakche et al., 2024). The cluster tests were applied to determine if the RSA coefficients, calculated across all epochs, were significantly different from zero, and if there was greater fidelity (larger RSA coefficients) in neural responses to subsequently recognized images versus those subsequently missed. The latter analysis involved repeating the RSA procedure separately on each participant’s two subsets of epochs (hits, misses) after equating trial numbers, then subtracting the RSA coefficient time courses for miss trials from those for hits. Two participants were excluded from this subtraction analysis after the RSA performed on their hits data resulted in negative coefficients during the first 300 ms post saccade, which, in the early visual and semantic representations, were more than two standard deviations below the group mean; a pattern likely attributable to noise in the estimates of dis-similarity from relatively small trial numbers.

#### 2.5.4 Relating saccade-alpha phase locking to neural representations and memory performance

Our primary hypotheses in this investigation were two-fold. First, that phase-locking one’s saccades to ongoing alpha oscillations improves the neural representation of the subsequently fixated visual stimulus; and second, that this improvement is related to better memory encoding of that visual stimulus. The analyses outlined in the previous sections focused on establishing the presence of the relevant subsequent memory effect in alpha ITPC and the emergence of visual and semantic representations in post-saccade neural responses. Once found, we could then test the primary hypotheses.

To determine whether phase-locking one’s saccades improves neural representation of fixated stimuli, we examined the relationship between pre-saccade alpha ITPC and the fidelity of the visual and semantic representations that emerged from those saccades on the encoding trials. We note that the best test of this hypothesis would involve analysis at the single trial level, where post-saccade RSA coefficients could be predicted from pre-saccade alpha phase. However, the operational definitions of the two constructs, using ITPC and RSA, required aggregating responses across trials. We thus turned to the next best alternative, which was to conduct the analysis at the group-level, or between subjects. Here, we predicted that those participants who showed greater saccade-alpha phase-locking would also show better fidelity visual and semantic representations, once their eyes landed on that stimulus. To test this, we computed Pearson correlation coefficients between participants’ pre-saccade alpha ITPC values, extracted from the 200 ms preceding the first saccade to each image, and the fidelity of visual and semantic representations, after the execution of those saccades. The 200 ms ITPC window was selected to encompass the alpha-band effect identified in the subsequent memory contrast. The fidelity values were extracted and averaged from a 100-200 ms window after the onset of the saccade, chosen to capture the peak data-model correspondence across the three examined levels (see Figure 3).

Next, to test whether alpha ITPC and visual and/or semantic representations were related to better overall memory performance on the subsequent recognition test, including the correct rejection of Foil images, we constructed a multiple regression model that attempted to predict each participants’ d-prime value from these properties. We extracted each participant’s mean pre-saccade alpha ITPC (-200 to 0 ms, before saccade onset) and the neural fidelity of early visual, late visual, and semantic representations (100-200 ms, after saccade onset), averaged across all encoding trials. We note that this is a rather coarse assessment of whether these properties explain variation in memory encoding, as it is done at the task-rather than the item-level, but one which, if successful, would provide strong evidence that saccade-alpha phase-locking and post-saccade neural representations are directly related to one’s overall ability to encode visual objects in memory.

### 2.6 MEG Source Localization

Finally, we re-examined visual- and semantic-representations that emerged after the first saccades onto each object during the encoding phase, in activity localized to the cortical surface using noise-normalized L2 minimum-norm estimation. Covariance matrices were calculated from the concatenation of 100 ms windows extracted from the central fixation period of each trial using an automated regularization method (Engemann & Gramfort, 2015), which selects the best estimator from three options: the empirical covariance, diagonal loading, and a data-driven extension of the Ledoit-Wolf shrinkage model (Ledoit & Wolf, 2004). The T1 anatomical MRIs were used to define cortical surface models via Freesurfer’s automated segmentation algorithms (recon-all; Dale, Fischl, & Sereno, 1999; Fischl, Sereno, & Dale, 1999; Fischl et al., 2004, Fischl, 2012). The MRI coordinate space for each participant’s anatomical T1 was co-registered with the MEG coordinate space based on the location of fiducials and the three head position indicator coils. Forward models were estimated for each participant’s cortical surface using a single-layer boundary element model. Following past work (Flick, Abdullah, & Pylkkänen, 2021), we set the inverse operator’s regularization signal-to-noise parameter to 2 and used “fixed” orientation sources that were constrained to lie perpendicular to the participant’s cortical surface (see also Dale & Sereno, 1993; Lin, Belliveau, Dale, & Hamalainen, 2006; Gwilliams, Lewis, & Marantz, 2016). Application of each participant’s inverse operator to their saccade-locked epochs yielded source-time courses, time-locked to the first eye movement to the image on each encoding trial.

Source-localized responses were examined in three bilateral regions of interest (ROIs) defined from the Desikan-Killiany automated cortical parcellation (Desikan et al., 2006) and motivated by previous literature. First, as past evidence has demonstrated that alpha oscillations related to visual processing and memory localize to superior parietal and lateral parieto-occipital areas (e.g., Tuladhar et al., 2007) we combined the atlas’s inferior and superior parietal parcels on the lateral surface into one parieto-occipital ROI. Second, we defined an ROI that encompassed Brodmann areas 17 and 18, shown by Devereux et al., (2018) to encode early visual representations, by combining the atlas’s parcels for the lingual gyrus and pericalcarine cortex. Inspection of the source-localized evoked responses to the image onsets and eye movements revealed early (e.g., 100 ms) response components in these two regions, but also in the neighbouring medial parietal cortex. We thus added to this ROI, two additional parcels to capture the spatial extent of the early visual response: the precuneus and cuneus. Third, to examine responses in the ventral temporal lobe that have been implicated in the semantic or conceptual representation of visual objects (Clarke et al., 2018; Devereux et al., 2018; von Seth et al., 2023), we defined the final ROI from the combination of the atlas’s parcels for the fusiform gyrus, inferior temporal gyrus, entorhinal cortex, and parahippocampal gyrus. For convenience, we refer to these regions, respectively, as the lateral parieto-occipital, medial parieto-occipital, and ventral temporal ROIs.

## 3. RESULTS

### 3.1 Behaviour

The average participant hit rate at recognition was 73.40% (SD = 12.12%), equivalent to 183 out of 250 correctly recognized images and 67 misses. This was significantly different from chance (at 50%), demonstrated by a non-parametric resampling test using 1000 random sign-flips of each participant’s deviation from chance-level (p < 0.001). The mean d-prime and corrected recognition values across participants were 1.64 (SD = 0.56) and 54.44% (SD = 16.17%), respectively. The rate of false alarms for novel items at recognition was relatively low, with a mean of 18.96% (SD = 12.48%) equivalent to incorrectly endorsing only 47 of 250 new items as old. On the basis of their behavioural results, two participants were excluded from the MEG analyses of subsequent memory performance. In both cases this was due to a hit rate above 90%, which resulted in low trial numbers after equating subsequent hit and miss trials.

### 3.2 Eye-tracking

Across all encoding trials, participants made an average of 5.40 (SD = 1.15) fixations on each image with a mean duration of 408.55 ms (SD = 124.70 ms) per fixation. Non-parametric resampling tests, using 1000 random flips of each participant’s hit/miss data, were used to assess the statistical significance of the eye movement difference related to subsequent memory. As shown in Figure 2, images that were later recognized were encoded with significantly more fixations (M_hit_ = 5.53, M_miss_ = 5.08; p < 0.001) and a corresponding reduction in the mean fixation duration (M_hit_ = 487.62 ms, M_miss_ = 546.44 ms, p < 0.001), matching past reports of associations between visual sampling and subsequent memory (Loftus, 1972; Chan et al., 2011; Olsen, et al., 2016). The difference in latency of the first fixation on the image after its appearance (i.e., reaction time) was trending toward statistical significance (p = 0.073), however the numerical difference between hits (M_hit_ = 253.00 ms) and misses (M_miss_ = 259.32 ms) was less than 7 milliseconds. There was no significant difference in the amplitudes of the initial saccades to images that were later recognized (M_hit_ = 8.18 dva) versus those that were not (M_miss_ = 8.20 dva; p = 0.569).

**Figure 2.**
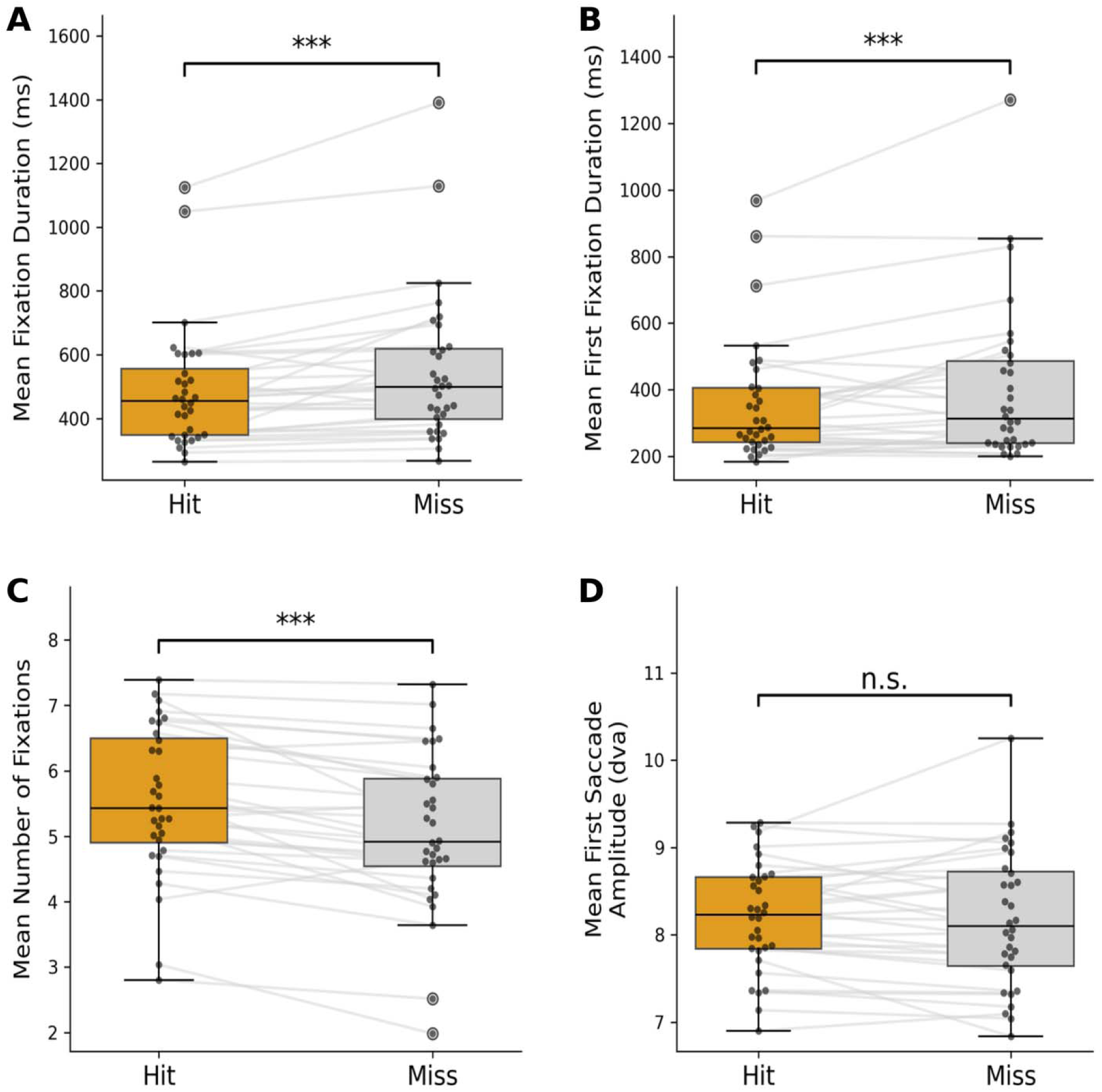
Eye-tracking results. (A) Mean fixation durations, (B) mean first fixation durations; (C) mean number of fixations, and (D) mean amplitude of the first saccade to the images, for subsequent Hits and Misses. *** indicates a statistically significance difference, corresponding to p < 0.001. n.s. indicates no significant difference.

The mean duration of the first fixation on the image across all trials was 373.24 ms (SD = 454.89 ms). This duration was significantly shorter when viewing images that were subsequently recognized (M_hit_ = 358.83 ms) relative to those that were not (M_miss_ = 402.28 ms, p < 0.001), demonstrating that the association between greater visual exploration and better subsequent memory was already present upon the first fixation on the image. This effect was not present during the pre-image interval, where there was no difference in the number of saccades executed during the presentation of the fixation square (M_hit_ = 2.95, M_miss_ = 2.92, p = 0.282). There was also no difference in the mean duration of the fixation that preceded the first saccade onto images that were later recognized (M_hit_ = 729.45 ms) versus those that were not (M_miss_ = 745.19 ms, p = 0.161), indicating that participants maintained central fixation for approximately the same amount of time in each case. The magnitude of these fixation durations (greater than 700 ms, on average) further demonstrated that participants adhered to the instruction that they return their gaze to and maintain fixation on the central square to start each trial. We also examined potential differences in eye movements related to the side of the screen on which the image was presented, but found no such evidence. There were no significant differences between the number of fixations (M_left_ = 5.43, M_right_ = 5.37; p = 0.235), their mean duration (M_left_ = 504.06 ms, M_right_ = 509.21 ms; p = 0.524), nor the mean duration of the first fixation on the image (M_left_ = 372.61 ms, M_right_ = 372.69 ms; p = 0.991) when comparing left and right presentation trials.

### 3.3 Grand Average MEG Sensor Responses

We began our analysis of the MEG data by examining grand average responses to the presentation of the image on each encoding trial, irrespective of subsequent memory. Figure 3 displays the average event-related field response (the evoked response) elicited by the image onset (A) and the onset of the first saccade to the image (B), as well as frequency-resolved time courses of power (C, D) and inter-trial phase coherence (E, F). In the time-domain, the image elicited prominent response peaks at approximately 100, 200 and 250 ms, followed by an extended response component that spanned approximately 300-500 ms. The topography of the latter component (outgoing fields over the left hemisphere, ingoing over the right) was a reversal from the earlier patterns at approximately 100 ms after image onset, but matched the topography emerging after the first saccade (Figure 3B). This is consistent with a recent report that saccades and image presentations elicit opposite polarity MEG responses (Amme et al., 2024). When considering the mean onset time of the first saccade to the image (approximately 250 ms) and the latency of the first visual peak following this saccade (approximately 100 ms later), this straightforwardly explains the extended late visual response to the image as emerging from saccade-locked neural activity.

**Figure 3.**
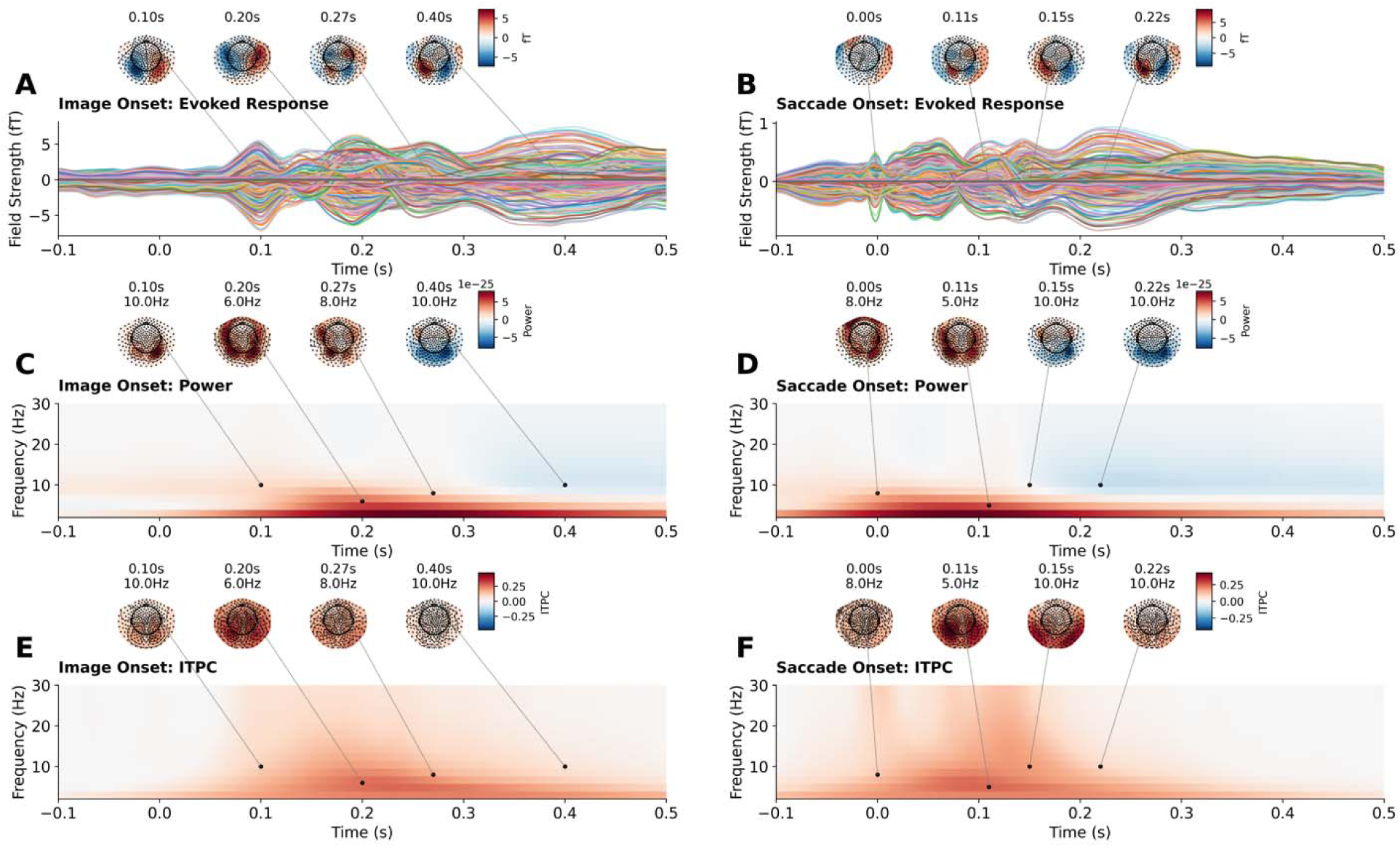
Grand average responses time-locked to the onsets of images (left column) and initial saccades (right column), regardless of subsequent memory. (A) Average time-domain response to the onset of the image on each encoding trial. Each colored line depicts the response from one MEG channel. (B) Average time-domain response to the onset of the initial saccade from central fixation to the image, on each encoding trial. (C) Power time-locked to image onset. (D) Power time-locked to initial saccade onset. (E) ITPC time-locked to image onset. (F) ITPC time-locked to initial saccade onset.

Inspection of the grand average power time course (Figure 3C) following image onsets revealed a peak at approximately 200 ms, which was predominantly constrained to the lower frequency ranges (5-12 Hz). In the alpha and beta bands, this was followed by a reduction in power beginning at approximately 300 ms, to levels below the pre-image baseline interval (i.e., event-related desynchronization). ITPC showed a similar peak at 200 ms, over parietal sensors, and a return toward baseline 300-500 ms following the image onset (Figure 3E).

These patterns were also evident in time-frequency analyses time-locked to the first saccade to the image (Figure 3D & F). Saccades were preceded by, and occurred in the midst of, heightened low frequency power and phase alignment over parieto-occipital sensors (Figure 3D & F). The power time course again showed a reduction in the alpha and beta frequency ranges (event-related desynchronization), which began approximately 150 ms after saccade onset, and coincided with a return toward baseline in ITPC. This is consistent with previous reports that saccades are followed by event-related desynchronization in the alpha and beta frequency ranges (Popov et al., 2021; Popov & Staudigl, 2023; Wu et al., 2025), and again suggests that late responses to the images, this time in the frequency domain, may be explained by neural activity associated with saccadic eye movements that bring the image into foveal vision. Importantly, and discussed in greater detail in the following sections, several follow-up analyses indicated that our memory-related effects in pre-saccade phase locking were not simply due to the transient increase in ITPC elicited by the onset of the image at the start of each trial.

### 3.4 Pre-saccade alpha ITPC predicts subsequent image memory

Past work demonstrated that successful encoding of visual scenes was associated with heightened ITPC in the alpha frequency band before saccade onsets, suggesting that saccades during successful encoding were executed at more consistent phases of ongoing alpha oscillations (Staudigl et al., 2017). Here, we asked whether the same subsequent memory effect was observable when participants executed a single eye movement from a central fixation square to a visual image presented in the periphery.

A cluster-based permutation test on the time course of alpha-band ITPC (8-12 Hz) at bilateral parietal sensors revealed significantly greater ITPC before saccades onto images that were subsequently recognized (hits) compared to images that were not (misses; -168 ms to 0 ms before saccade onset, cluster p-value = 0.003). As shown in Figure 4, visualization of the saccade-locked difference in ITPC revealed a pattern centered over parietal and occipital MEG channels. When we expanded the time window of analysis into the post-saccade interval (300 ms pre-saccade to 300 ms post-saccade), the significant alpha-band cluster spanned from 168 ms before the saccade to 110 ms afterward (cluster p-value < 0.001). The peak alpha ITPC difference between subsequent hits and misses, including the post-saccade interval, was approximately 60 ms before saccade onset.

**Figure 4.**
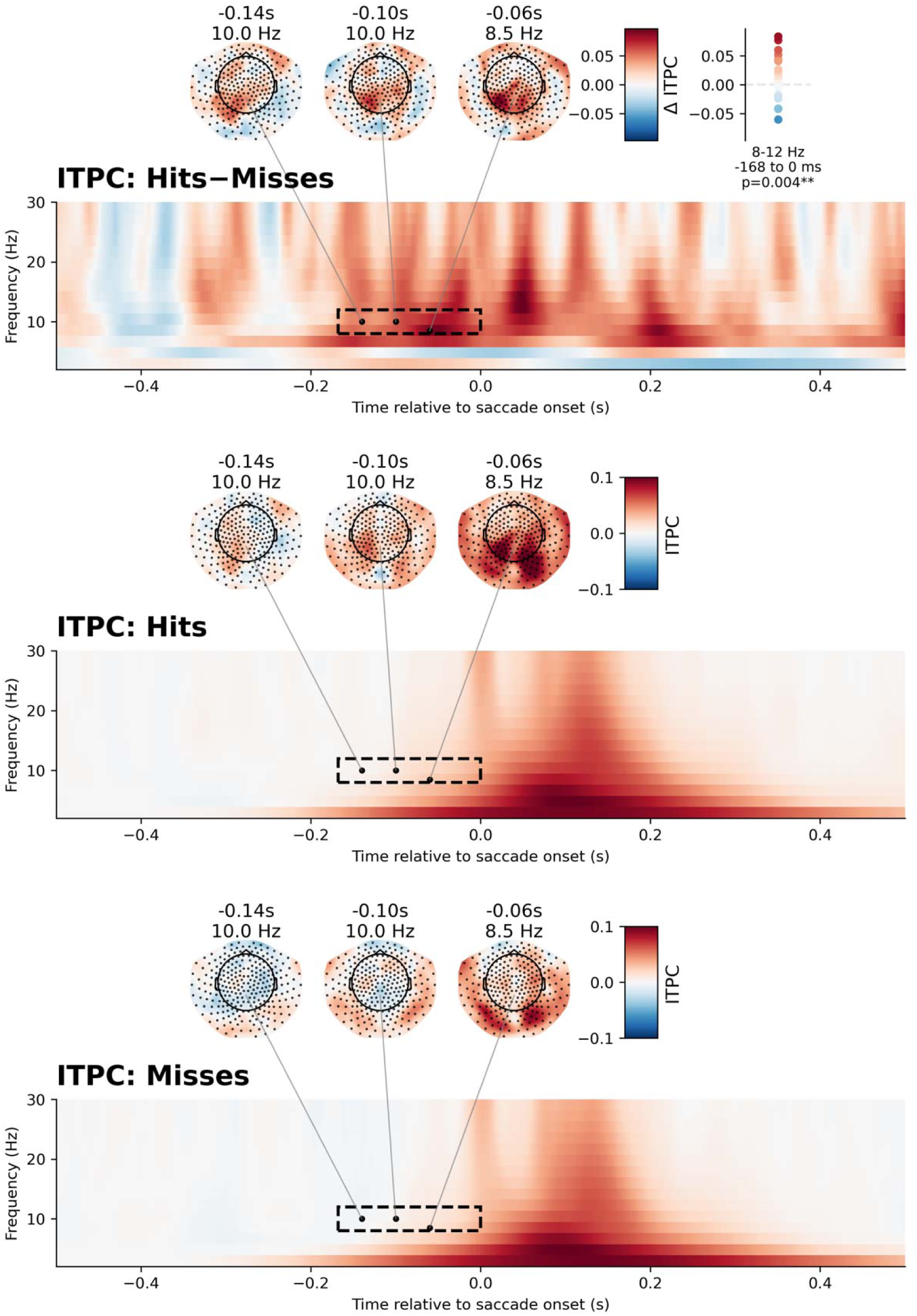
Spectrograms of ITPC values time-locked to the onset of the first saccade (0.0s) to the images during encoding. Each spectrogram was extracted and averaged responses from all MEG channels, visualized in the topographic channel maps. The significant alpha-band (8-12 Hz) cluster at parietal channels, highlighted in the black rectangle, contained greater ITPC preceding saccades onto subsequent hits, from -168 ms to 0 ms before saccade onset. Mean alpha ITPC difference values from each participant, in this time window, are displayed in the top-right corner. The bottom two panels display the spectrograms of ITPC, relative to a baseline during central fixation, time-locked to the first saccades to subsequent hits and subsequent misses.

We conducted several follow-up analyses of the ITPC subsequent memory effect. First, to test whether the pre-saccade effect was an artifact of temporal smearing during wavelet convolution, stemming from activity at or after onset of the eye movement (e.g., the saccadic spike potential; Thickbroom & Mastaglia, 1985), we cropped the saccade-locked epochs at increasingly early latencies before the saccade (-10, -20, -30, … -90 ms), prior to the convolution step, and performed one-tailed t-tests on the mean alpha ITPC difference, extracted at 8, 10, and 12 Hz. These tests revealed that the mean ITPC difference from 167 ms before the saccade (the original cluster boundary), up to the end of the cropped epoch (e.g., -167 to -10 ms; -167 to -20 ms, etc.), remained statistically significant even when the epochs were cropped 100 ms before the saccade was executed, indicating that the memory-related difference was not an artifact of activity at or after saccade onset. We also note that there was no difference in saccade amplitude between subsequent hits and misses (see Section 3.2), suggesting that the ITPC effect was not due to differences in preparatory muscle activity.

Second, because differences in ITPC might stem from underlying differences in power – which can cause differences in the signal-to-noise ratio of phase estimates (van Diepen & Mazaheri, 2018) – we performed additional tests for these power differences. Although there was numerically greater alpha power prior to saccades to subsequent hits (see Supplementary Figure 1), no clusters were found in the comparison of pre-saccade alpha power time courses, and mean alpha ITPC and power were not correlated across participants in the spatiotemporal window where the ITPC effect was observed (Pearson correlation: r = 0.082, p = 0.668), contrary to what would be expected if power was driving the effect. Next, since pre-saccade alpha phase differences might be attributable to visual responses elicited by short previous fixations (Nikolaev et al., 2013, 2016), we repeated our analysis with an additional inclusion criterion: that the preceding fixation on the central square be at least 400 ms in duration. We again found significantly greater alpha-band ITPC before saccades onto subsequent hits compared to subsequent misses at the parietal sensors (p = 0.030), replicating the original result. These follow-up analyses bolster the notion that the memory-related difference in alpha ITPC is a genuine difference in alpha phase alignment across trials.

Finally, we investigated whether the memory-related difference in pre-saccade alpha phase alignment was also present when examining later saccades, executed within the image itself. This would more straightforwardly match the result of Staudigl et al., (2017), who examined multiple saccades during the encoding of more complex visual scenes. Although there was numerically greater alpha-band ITPC before saccades within later remembered images, cluster-based permutation tests performed on parietal alpha ITPC failed to find statistically significant differences when time-locking to the 2nd and 3rd saccade on each encoding trial (all p-values > 0.098). Based on this result, we focused all the following analyses solely on the ITPC preceding the first saccade to the image.

In summary, the first saccades onto images that were later remembered were preceded by greater ITPC in the alpha frequency band, relative to saccades onto images that were later forgotten. Follow-up tests indicated that the pre-saccade effect was not explained by differences in responses elicited by previous fixations, by frequency-domain artifacts of preparatory muscle activity (i.e., the saccadic spike potential), or by underlying differences in signal to noise ratios (i.e., power) when estimating phase angles. Finally, the effect was specific to the first saccade onto the image, executed from central fixation, suggesting that the memory-related difference is unique to an image arriving in foveal vision for the first time. This finding sets the stage for testing our primary hypothesis that greater saccade-alpha phase-locking leads to improved neural representations of the then-foveated visual stimulus, supporting memory encoding.

### 3.5 Saccade-locked Representational Similarity Analysis

Following past studies (e.g., Clarke et al., 2018), we used RSA to examine the emergence of visual and semantic representations in distributed patterns of neural responses. We note that while several previous investigations have examined the time course of these representations time-locked to the onset of a visual stimulus, relatively few have examined how these representations emerge when time-locking to saccades (e.g., Auerbach-Ausch & Deouell, 2024; Fakche et al., 2024).

Figure 5 displays the time-courses of model-data similarities, or RSA coefficients, for each level of representation measured at the MEG channels. When time-locking to the onset of the image, statistically significant evidence of early (Fig 5A; cluster: 137-457 ms, cluster p < 0.001) and late (Fig 5D; cluster: 256-460 ms, cluster p < 0.001) visual representations was found in the MEG responses, while there was no evidence for the presence of semantic representations (Fig 5G, cluster p > 0.05). When time-locking to the onset of the first saccade to the image, we found statistically significant evidence of early visual (Fig 5B, cluster: 0-323 ms, cluster p < 0.001), late visual (Fig 5E, cluster: 80-311 ms, cluster p < 0.001), and semantic (Fig 5H, cluster: 109-153 ms, cluster p = 0.047) representations in MEG-measured responses. Inspection of the RSA coefficient time course suggested that visual representations may also have been present prior to saccade execution, matching past results that have demonstrated parafoveal extraction of visual information about upcoming saccade targets (Fakche et al., 2024). This was confirmed in a follow-up test, which revealed significant early visual representations (Fig 5B, -85 to -37 ms, cluster p = 0.022) in the 300 ms before the onset of the eye movement. No significant late visual or semantic representations were found prior to saccade onset.

**Figure 5.**
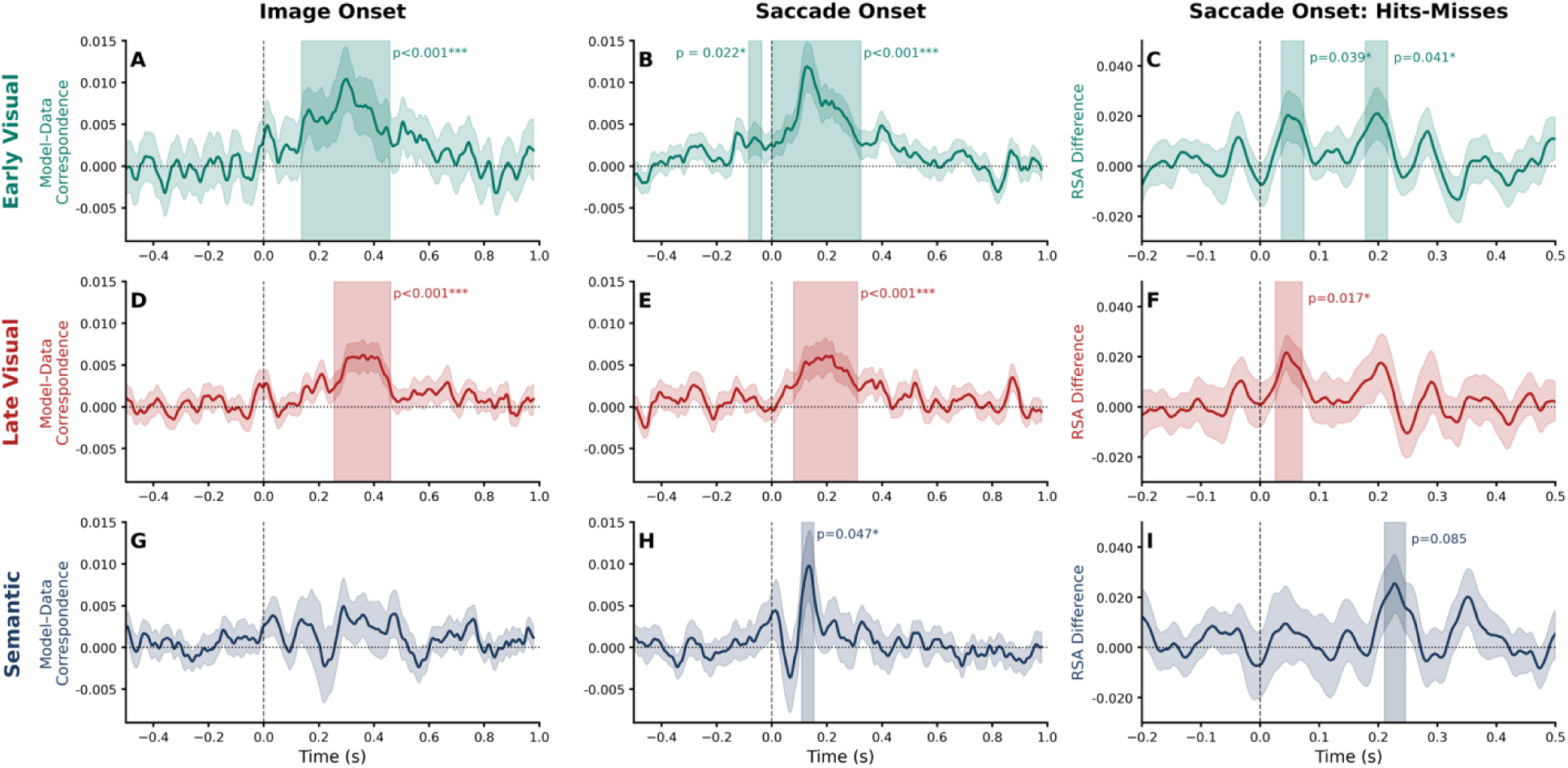
Time courses of RSA coefficients for the early visual (top-row), late visual (middle row) and semantic (bottom row) representations. Sub-plots show the time-course of representations relative to onset of the image (A, D, G) and the first saccade to the image (B, E, H), separately for early visual (A-B), late visual (D-E), and Semantic (G-H). The right-most column displays the difference in the fidelity of early visual (C), late visual (F) and semantic (I) representations, time-locked to the first saccade onto the image, subtracting misses from hits. Shaded regions indicate temporal clusters, with statistical significance indicated by the cluster p-values.

We also examined group-level differences in the fidelity of visual and semantic representations related to subsequent memory. For each participant, we subtracted the time course of RSA coefficients calculated from the set of images that were later missed, from those that were later recognized. We then performed cluster-based permutation tests against zero on the fidelity differences from 0-300 ms after the first saccade onset. We found transiently but significantly greater fidelity of early (Fig 5C, clusters: 36-74 ms, cluster p = 0.039) and late (Fig 5F, cluster: 26-71 ms, cluster p = 0.017) visual representations after saccades onto subsequently recognized images, compared to those that were subsequently missed. The early nature of these effects, beginning within 30 ms of the saccade onset, suggests they may have been mediated by parafoveal processing of the image, although there was also a second cluster that displayed greater fidelity in early visual representations later in time (178-216 ms, cluster p = 0.041). Greater semantic fidelity was also observed for subsequent hits compared to subsequent misses later in time, but this effect was above the threshold of statistical significance (Fig 5I, cluster: 211-246 ms, cluster p = 0.085).

### 3.6 Pre-saccade alpha phase locking predicts visual representational fidelity

Having established the presence of (a) pre-saccade alpha phase-locking differences related to subsequent memory (Figure 4) and (b) evidence of visual and semantic representations in post-saccade neural activity, whose fidelities also relate to subsequent memory (Figure 5), we tested the relationship between the two. Based on our hypothesis that phase-locking one’s saccades improves post-saccade neural representations, we predicted that those participants who show greater alpha ITPC prior to the first saccade to the image on each trial, would also show better neural representations of those images, once the eyes landed upon them.

Consistent with this hypothesis, and shown in Figure 6, pre-saccade alpha ITPC, extracted from the parietal channels during the 200 ms before saccade onset, was positively correlated with the fidelity of early visual (100-200 ms: r = 0.46, p = 0.010) representations that emerged in neural responses after those saccades. A slightly weaker positive correlation was found between pre-saccade alpha ITPC and the fidelity of late visual representations (r = 0.32, p = 0.081). Although we chose the 100-200 ms post-saccade window of the RSA coefficients to capture the overlapping peaks across all three levels of representation, post-hoc inspection of an earlier window (50-150 ms post-saccade) revealed even stronger correlations between alpha ITPC and the fidelity of visual representations (early visual, 50-150 ms: r = 0.50, p = 0.005; late visual, 50-150 ms: r = 0.33, p = 0.077). No significant relationship was found between alpha ITPC and the fidelity of post-saccade semantic representations in the MEG sensor data (p > 0.50). Likewise, no significant relationships were found between pre-saccade alpha power, extracted from the 200 ms window before onset, and any of the three levels of representation (maximum correlation coefficient = 0.13, all p-values > 0.47). This demonstrates that those participants who executed saccades at consistent alpha phases, also showed better visual-based representations of the images in neural activity, once those saccades were executed.

**Figure 6.**
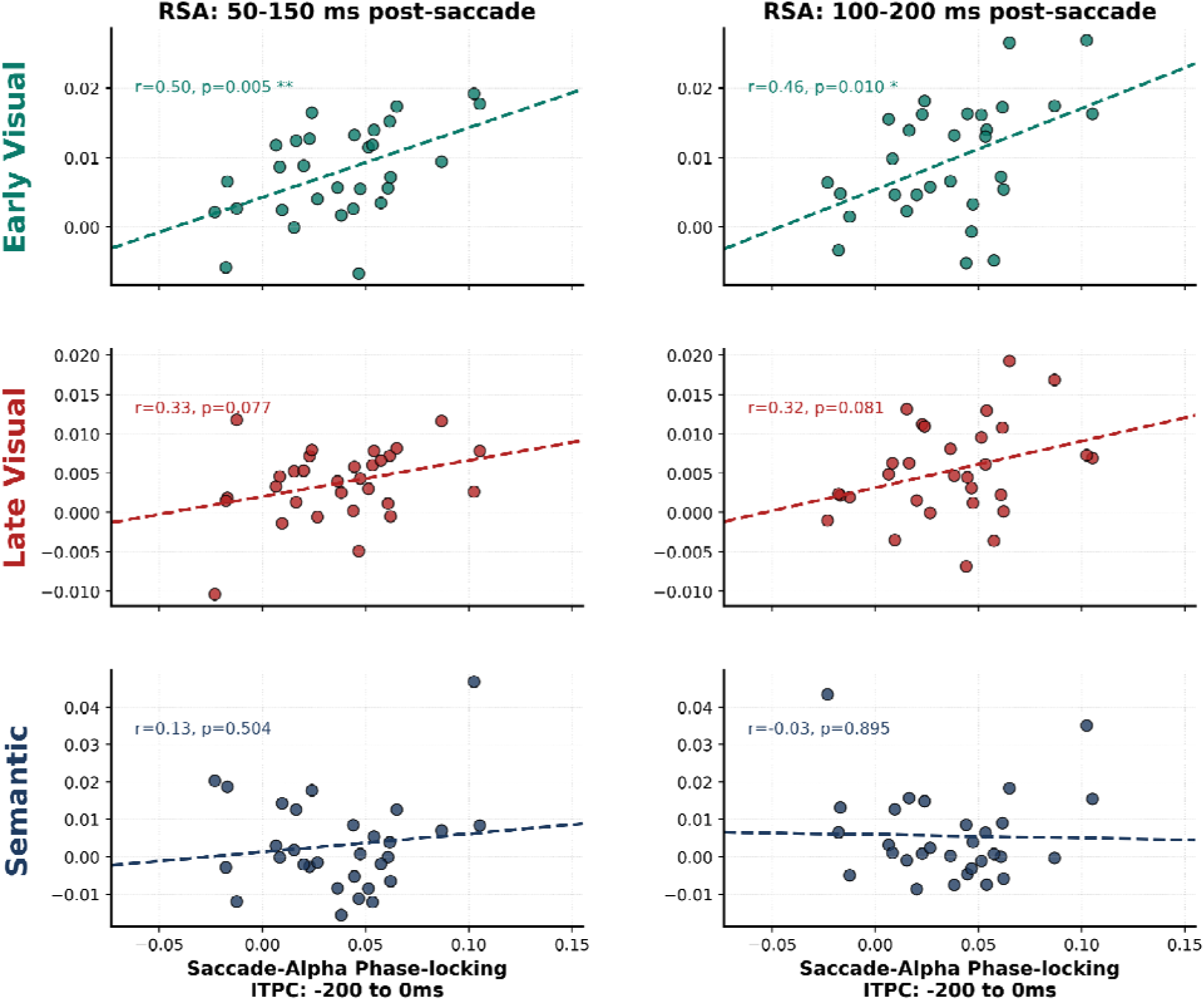
The relationship between the phase-locking of saccades to alpha oscillations and the fidelity of early visual (top-row), late visual (middle-row) and semantic (bottom-row) representations that emerge after those saccades, calculated across all sensors. RSA coefficients were extracted and averaged across two overlapping windows: 50 to 150 ms and 100-200 ms after saccade onset. Saccade-alpha phase-locking was calculated as the mean alpha-band ITPC at the parietal sensors in the 200 ms before saccade onset, relative to a central fixation interval.

The second piece of our hypothesis concerned the relationship with subsequent memory. We posited that the improved neural representations that result from phase-locking one’s saccades to alpha oscillations – as evidenced above – would facilitate improved memory encoding. We note that evidence for this was provided by the pair of results described earlier: (i) that the fidelity of visual representations was higher when the eyes landed on images that were subsequently remembered, relative to those subsequently forgotten (Figure 5); and (ii) that greater saccade-alpha phase-locking was associated with improvements in those fidelities (Figure 6).

To further test this connection between greater saccade-alpha phase-locking, fidelity of representations and subsequent recognition memory, we constructed a multiple regression model that attempted to explain variance in overall memory performance, operationalized using d-prime values, from each participant’s pre-saccade alpha ITPC, and the fidelity of their post-saccade representations (extracted from 100-200 ms, post-saccade). In other words, this analysis attempted to explain participants’ overall recognition memory in the experiment, including their ability to correctly reject novel items at test, as a function of 3-4 properties from the first saccade to the images at encoding, averaged across all trials.

The fully specified model, containing each participant’s mean pre-saccade alpha ITPC and all three representational fidelities, did not reach the threshold for statistical significance but was trending in that direction (Multiple R^2^ = 0.247, F_4,27_ = 2.208, p = 0.095). Interestingly, the only coefficient that was significantly different from zero was that of the semantic representation 100-200 ms after saccade onset (Estimate = 21.120, SE = 9.578, t = 2.212, p = 0.0356). A post-hoc comparison between a model that included only the pre-saccade alpha ITPC, and one that also included the post-saccade semantic fidelity, revealed a significant improvement in model performance when each participant’s mean semantic representational fidelity was added to the model (Multiple R^2^ for model_0_ = 0.073, Multiple R^2^ for model_1_ = 0.199, F_1,29_ = 4.591, p = 0.041). Adding either the early or late visual fidelity did not significantly improve the models’ abilities to explain variance in the overall memory scores (both p-values > 0.300). Overall, these results indicate that while phase-locking one’s saccades to alpha oscillations improves perceptual (i.e., visual) representations, and these two properties distinguish images that will or will not be subsequently recognized (Figures 5 & 6), the ability to rapidly extract semantic representations of an image, once the eyes land on it for the first time, may be a better predictor of participants’ overall performance on the current subsequent memory task, operationalized in d-prime values.

### 3.8 Localization of saccade-locked visual and semantic representations

Finally, we used source localization to define the areas of the brain showing evidence of visual and semantic representations that emerged in neural activity after the first saccade onto the images. Saccade-locked MEG epochs were projected to each participant’s cortical surface using noise-normalized L2 minimum norm estimation. We then repeated the sliding window RSA analysis on source time courses localized to three bilateral regions of interest (ROIs): medial parieto-occipital, lateral parieto-occipital, and the ventral temporal lobes (see Section 2.6 for ROI definitions). For each RSA, model-derived similarities were compared to the vectors of source activity across all constituent sources in the ROI, in sliding 50 ms time windows centered on every individual time point. Cluster-based permutation tests assessed whether the coefficients were significantly greater than 0, in a 0-500 ms analysis window after saccade onset.

As shown in Figure 7, cluster-based permutation tests revealed that early visual representations were present in all six ROIs. In the lateral and medial parieto-occipital regions, evidence of early visual representations appeared immediately upon saccade onset (left medial: 0-452 ms, cluster p < 0.001; right medial: 0-343 ms, cluster p < 0.001; left lateral: 0-352 ms, cluster p < 0.001; right lateral: 0-431 ms, cluster p < 0.001) whereas significant clusters emerged in the ventral temporal ROIs by approximately 100 ms post-saccade (left ventral temporal: 98-194 ms, cluster p = 0.006; right ventral temporal: 100-247 ms, cluster p < 0.001). Follow-up tests further confirmed that signals reflecting early visual representations emerged in all four parieto-occipital ROIs prior to saccade onset (onset time of pre-saccade clusters: left medial: -135 ms; right medial: -152 ms; left lateral: -119 ms; right lateral: -123 ms). The fidelity of the early visual representations was graded across ROIs, with the greatest fidelity found in the medial parieto-occipital ROIs, followed by the lateral parieto-occipital ROIs and then the ventral temporal lobes.

**Figure 7.**
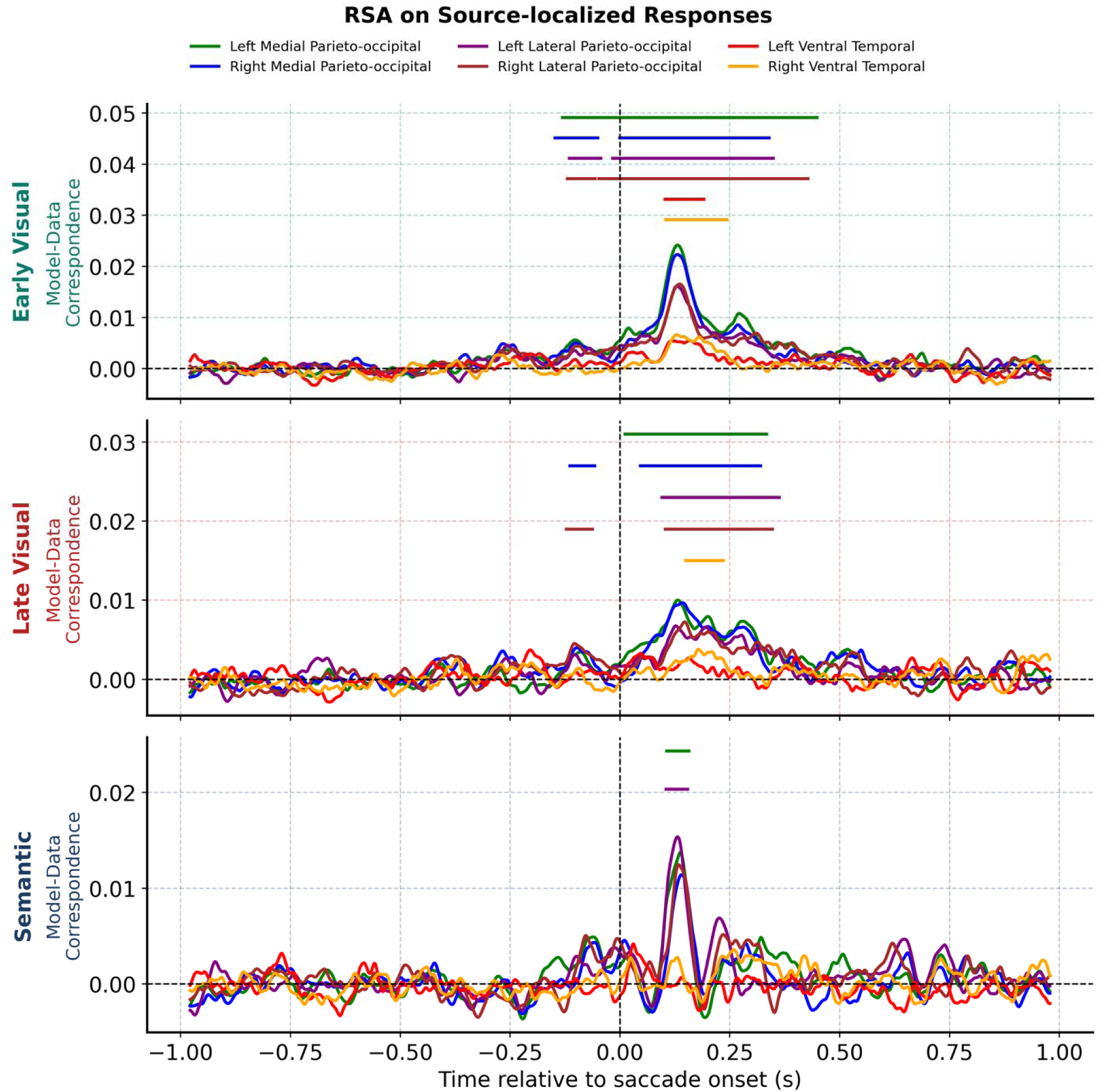
Representational similarity analysis performed on saccade-locked responses localized to six regions of interest using the early visual (top-row), late visual (middle-row) and semantic (bottom-row) models. Horizontal lines indicate significant temporal clusters (p < 0.05) of RSA coefficients that were greater than zero in the corresponding region.

Evidence of late visual representations were also found in the medial parieto-occipital (left: 8-337 ms, cluster p < 0.001; right: 43-323 ms, cluster p < 0.001) and lateral parieto-occipital ROIs (left: 92-366 ms, cluster p < 0.001; right: 99-350 ms, cluster p < 0.001), as well as the right (146-239 ms, cluster p = 0.003), but not the left ventral temporal lobe. In the right medial (-117ms) and lateral parieto-occipital cortex (-126 ms), signals consistent with late visual representations also emerged before saccade onset. Resembling the pattern observed in the sensor-level RSA, there was a transient signal reflecting the semantic representation of the images that emerged in the left medial (103-160 ms, p = 0.029) and lateral parieto-occipital (101-157 ms, cluster p= 0.034) cortex after saccade onset. No significant evidence of semantic representations was found in the ventral temporal lobes at the group level.

In summary, the source-localized RSA revealed that signals consistent with visual representations of the peripheral images were present in parieto-occipital cortex at the onset of the saccade to those images, and were maintained throughout the following 400 ms. Saccade-locked visual representations were also present in the bilateral ventral temporal lobes, however these emerged approximately 100 ms after the onset of the eye movement that brought the image into foveal vision. Semantic representations of the images, defined from feature production norms, were present in the left parieto-occipital cortex for a brief window after saccade onset, suggesting that this correspondence might be driven by semantic properties that co-vary with visual information (e.g., “is red” as a feature of a stop sign).

Having identified the source-level representations of the foveated images, we returned to the question of whether phase-locking one’s saccades to alpha oscillations improves these representations. We repeated the between-participant analysis of the relationship between pre-saccade alpha ITPC and the fidelity of the post-saccade representations, this time extracting the fidelity values from the source-estimated RSA data in each ROI. Here, instead of focusing on just a single time window, we extracted and averaged each participant’s RSA coefficients from each ROI, in five 100 ms windows after saccade onset starting from 0 ms and increasing in 50 ms increments. This provided a more comprehensive, spatiotemporal examination of how phase-locking one’s saccades might improve localized neuronal representation.

After correcting for multiple comparisons across the six ROIs and five time windows, there were no correlations between pre-saccade alpha ITPC and post-saccade representations that reached the level of statistical significance (corrected p < 0.05, using the False Discovery Rate). This is not a surprise, given the number of statistical comparisons considered in the correction. There were, however, notable trends toward significance, several of which matched the patterns observed in the sensor-level data. The strongest associations between saccade phase-locking and representational fidelity were found in left hemisphere ROIs, and predominantly in the early visual and semantic representations. Figure 8 visualizes these relationships, with the full set of results shown in Supplementary Figures 2 and 3.

**Figure 8.**
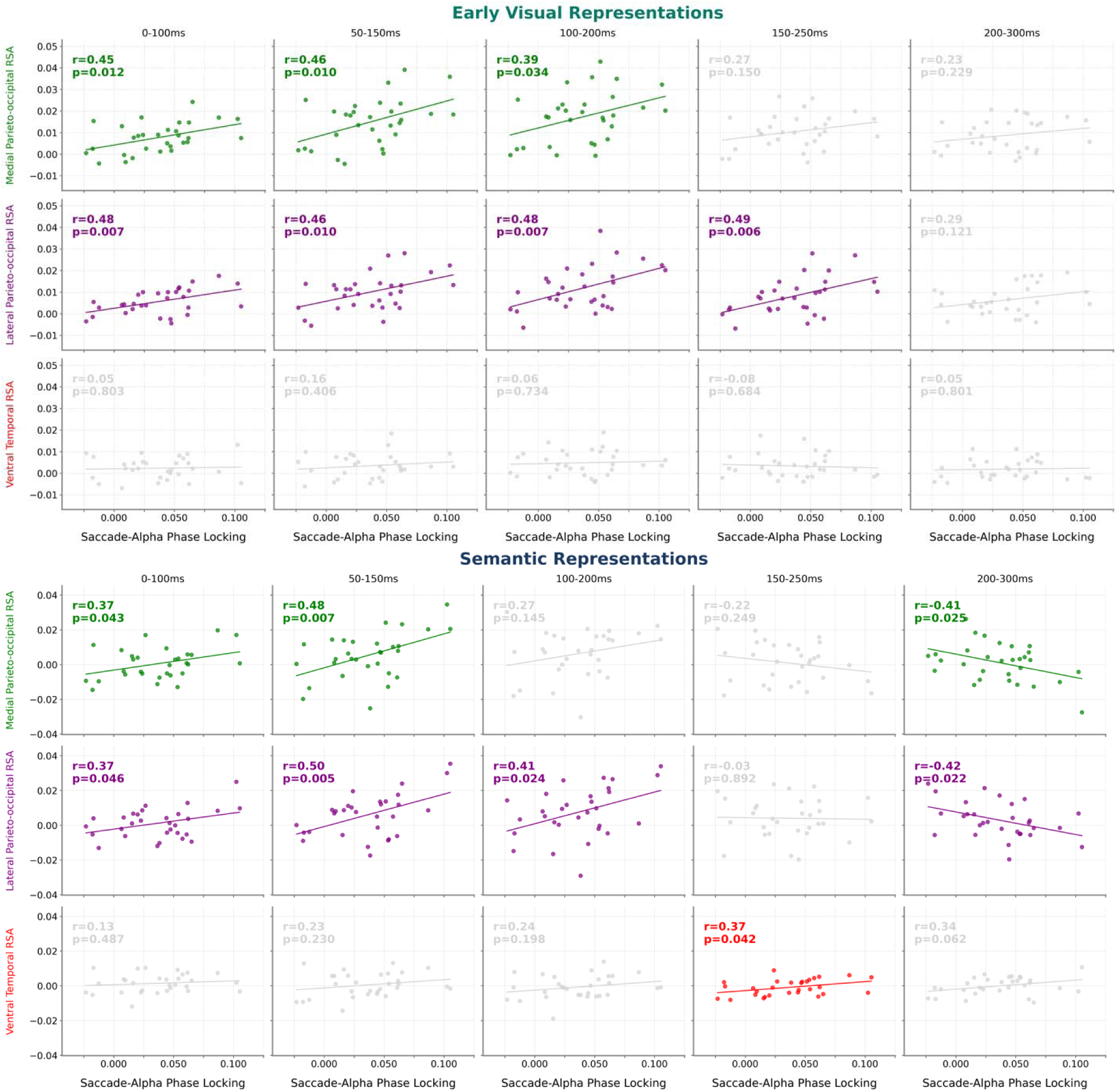
The relationship between saccade-alpha phase-locking and post-saccade representations in left hemisphere ROIs. Positive associations with the fidelity of early visual (top) and semantic (bottom) representations were found in the left lateral and medial parieto-occipital ROIs in the first 200 ms after saccade onset. The fidelity of semantic representations also showed a weak positive association with phase-locking in the left ventral temporal lobe from 150-250 ms after saccade onset, overlapping with the onset of negative associations in the parieto-occipital ROIs from 200 to 300 ms. Coloured scatterplots indicate Pearson correlation coefficients corresponding to an uncorrected p < 0.05.

Positive associations between saccade phase-locking and early visual and semantic representations emerged in left hemisphere parieto-occipital ROIs within 100 ms of saccade onset, followed by weaker positive correlations with semantic representations in the left ventral temporal lobe later in time (150-250 ms). Both associations indicate that those participants who more consistently phase-locked their eye movements to alpha oscillations showed better neural representation of the fixated images in these regions and at these time points. Interestingly, the positive association between phase-locking and semantic representations appeared in the left ventral temporal lobe during the same time window when significant visual representations of the fixated items were found in this region (97-198 ms; Figure 7), suggesting that saccade phase-locking may support the parallel emergence of semantic representations alongside those visual representations. Additionally, the positive association between phase-locking and semantic representation in the ventral temporal lobe overlapped with a switch to a negative association between pre-saccade ITPC and semantic representations in the parieto-occipital ROIs (200-300 ms), where participants with lower phase locking displayed better semantic representations in this later time window.

Although these results should be interpreted conservatively, given their secondary nature and the number of statistical tests performed, the pattern is consistent with a graded emergence of visual-to-semantic representations after fixation onset, which is facilitated when saccades are phase-locked with ongoing alpha oscillations. Finally, the negative associations later in time might suggest that when saccades are poorly synchronized with alpha oscillations, there is a delay in the visuo-semantic processing of the newly foveated visual stimulus.

## 4. DISCUSSION

The current results provide new evidence that phase-locking saccades to 8-12 Hz alpha oscillations, recorded over posterior brain areas, contributes to memory encoding by improving the neuronal representation of visual objects brought into foveal vision at the terminal of those saccades. The relationship between phase-locking and neural representation was found in activity localized to the left parieto-occipital cortex and ventral temporal lobe. Moreover, memory-related differences in how well neural activity represented visual objects were evident from the onset of the first saccades that brought those objects into foveal vision, suggesting that parafoveal processing may influence whether one remembers seeing an object later in time. We discuss each of these results, and their implications for our understanding of visual memory formation, in greater detail below.

### 4.1 Saccade phase-locking supports the neural representation of foveated visual objects

Our hypothesis that saccade phase-locking would lead to improved neural representation was motivated by accounts characterizing alpha oscillations as pulses of inhibition, wherein stimuli arriving at inhibitory phases result in dampened neural responses, relative to those that arrive outside of the inhibitory phase (Matthewson et al., 2011; Jensen et al., 2014; Gips et al., 2016). If this characterization is generalizable to active vision, and there is evidence suggesting that it is (Jensen et al., 2021), then eye movement timing must be coordinated so that foveal input does not arrive at a phase that results in suppression by the local alpha oscillation.

Following from this, we reasoned that the results of Staudigl et al., (2017) – showing that when saccades were phase-locked, there was improved scene encoding – might be partially explained by input arriving at the optimal phase. The improvement in neuronal processing, in turn, would result in better representation and cortical relay of the constituent elements fixated in a visual scene, resulting in a more accurate memory trace.

The current results validate this reasoning. Participants who kept their initial saccades tightly phase-locked to alpha oscillations showed better neural representations of the objects that they fixated on each encoding trial, and these two properties were each associated with better subsequent memory for individual objects. We defined better neural representations as greater correspondence between neural- and model-similarity, across encoding trials, in representational similarity analysis. In the case of visual representations, model-based similarity was defined by extracting and comparing the activation weights of a convolutional neural network model, whereas model-based semantic similarity was defined using semantic property norms. Past results have shown that these forms of representation capture meaningful properties of real-world concepts, and explain variance in sensory areas of the brain, in response to images of those concepts (Cree, McNorgan & McRae, 2006; Devereux et al., 2018; Clarke et al., 2018). We thus reason that when saccades are phase-locked to alpha oscillations, the resulting neuronal representations are truer to the physical form of the foveated content, and more accurately encode the denoted concept. In contexts where multiple items are fixated (e.g., visual scenes) the representations are then bound together by the hippocampal memory systems that synchronize with visual alpha oscillations (Olsen et al., 2012, 2013; Kragel et al., 2021) to form an improved relational memory trace.

While we did not find a strong relationship between participants’ phase-locking and visual RSA coefficients, and their overall memory performance (d-prime), we expect that this relationship might emerge when examining multiple saccades across multiple objects in a stimulus; e.g., predicting participants’ overall recognition memory for natural scenes from their average within-scene phase-locking and how well their post-saccade responses represented elements within those scenes. In addition to simply providing more data per trial, saccade phase-locking in this context might provide a means to capture how efficiently a viewer can encode a given scene in a constrained time window. This is a property that, we posit, is more likely to explain variance in a high-level memory outcome variable, such as d-prime, compared to the same properties extracted from just a single saccade to an isolated image. As will be discussed below, this might stem from a relationship between saccade alpha phase-locking and parafoveal processing, which ultimately supports greater visual sampling.

### 4.2 The role of parafoveal processing in memory encoding

Two current findings suggest that the degree to which one can parafoveally process an upcoming saccade target influences the likelihood that the visual stimulus at that target will be remembered later in time. First, in the contrast of subsequently remembered and forgotten images, time-locked to the first saccade onto those images, we observed very early (∼30 ms) differences in the fidelity of the neural representations, which were better for those later remembered. This was followed by slightly later (∼180 ms) differences, which also showed better neural representation, at the early visual and semantic levels, for subsequent hits (Figure 5). Second, pre-saccade alpha ITPC was positively associated with evidence for neural representation within the first 100 ms of the first saccade to objects (Figure 8).

We posit these findings are due, in part, to parafoveal processing. The average saccade latency after the image’s appearance was approximately 250 ms, and the results from the RSA collapsed across all trials (Figure 5 and 7) indicate that visual representations were present in neural activity prior to the onset of eye movements. We note that we cannot distinguish whether the later recognized images had properties that enabled parafoveal extraction, or if participants more actively engaged in an extraction process (i.e., a passive versus active explanation). In either case, the extraction of information prior to the saccade could explain the subsequent difference in fixation durations, once the eyes landed on the image. Images that were remembered were encoded with shorter first fixation durations (means: 358 vs. 402 ms) and, on average, 0.45 more fixations. Thus, the initial parafoveal processing might have led participants to spend less time fixating the image at their initial landing site and promoted the execution of more fixations overall, creating the attested memory encoding benefit that comes with greater visual sampling (Loftus, 1972; Chan et al., 2011; Olsen, et al., 2016).

What then, explains the early (0-100 ms) positive relationship between saccade-alpha phase-locking, and the fidelity of representations in parieto-occipital cortex, across participants? While some of this might be attributable to foveal information received once the saccade ended, due to the benefit of phase-locking saccades, we believe that parafoveal processing also has a role. Going back to accounts of alpha oscillations as pulses of inhibition (Jensen et al., 2021), one possibility is that some participants were able to release the representation of the peripheral image from alpha’s inhibition earlier in the oscillation, enabling attention to shift earlier and facilitate increased parafoveal processing, before saccade programming was completed and the eyes moved. This explanation would also account for the relationship with pre-saccade alpha ITPC in parieto-occipital cortex, as saccade target selection also depends on the phase-locked release of the peripheral representation from alpha inhibition. Those participants who were able to shift attention to the peripheral images earlier, prior to executing the saccade, might also have been able to better (i.e. more regularly) align their saccade onsets to the optimal alpha phase.

Finally, we note that our current results are unable to precisely tease apart the contribution of the image onset, which realigns alpha phases (Figure 2), and saccade phase-locking that takes place in addition to this. The prior work of Staudigl and colleagues (2017; and see Drewes & VanRullen, 2011; Pan et al., 2023), which used a design whereby saccades were executed within a scene, suggests that the same pattern would hold if the peripheral image was already on the screen well before the saccade was executed. We also have strong reason to believe that our results are not due to the evoked response. First, there was no significant memory effect in alpha power prior to saccade onset, which would be expected if the results were driven by the power increase elicited by the image’s appearance. Second, pre-saccade alpha power did not correlate with how well the images were represented in neural activity, once the eyes landed on them; that relationship was unique to the pre-saccade consistency of alpha phase. A natural avenue for future work will be to test the relationship between pre-saccade alpha phase and post-saccade neural representation in natural scenes or grids of visual objects; we would predict that when saccades are phase-locked, there would be improved neural representation of the constituent elements, fixated after those saccades.

### 4.3 Conclusion

The formation of visual episodic memories begins with the intake and neural processing of discrete visual samples, provided by saccadic eye movements. The current results demonstrate that when saccades are phase-locked to alpha oscillations, there is improved neural representation of the visual stimuli brought into foveal vision. This finding provides new mechanistic insight into how visual experiences are transformed into lasting memories. It indicates that precise coordination of saccade timing, with respect to neuronal oscillations, promotes more veridical neural representation of the resulting visual samples, from which accurate episodic memories can be constructed.

## Acknowledgements

We thank Korey Miller-Boyle, Ramsha Mahmood, Kinkini Monaragala, Hongyu Luo, and Rachel Goldfield for their valuable contributions to data collection. G.F. was supported by the Data Sciences Institute at the University of Toronto and by a Natural Sciences and Engineering Research Council of Canada (NSERC) Postdoctoral Fellowship. J.M. was supported by the Canada Research Chairs program and by NSERC grant RGPIN-2019-06515. J.D.R. received support as the Anne and Max Tanenbaum Chair of Baycrest Hospital and the University of Toronto, and from NSERC grant RGPIN-2018-06399. R.K.O. was supported by NSERC Discovery Award RGPIN-2024-06002.

## SUPPLEMENTARY MATERIALS

**Supplementary Figure 1.**
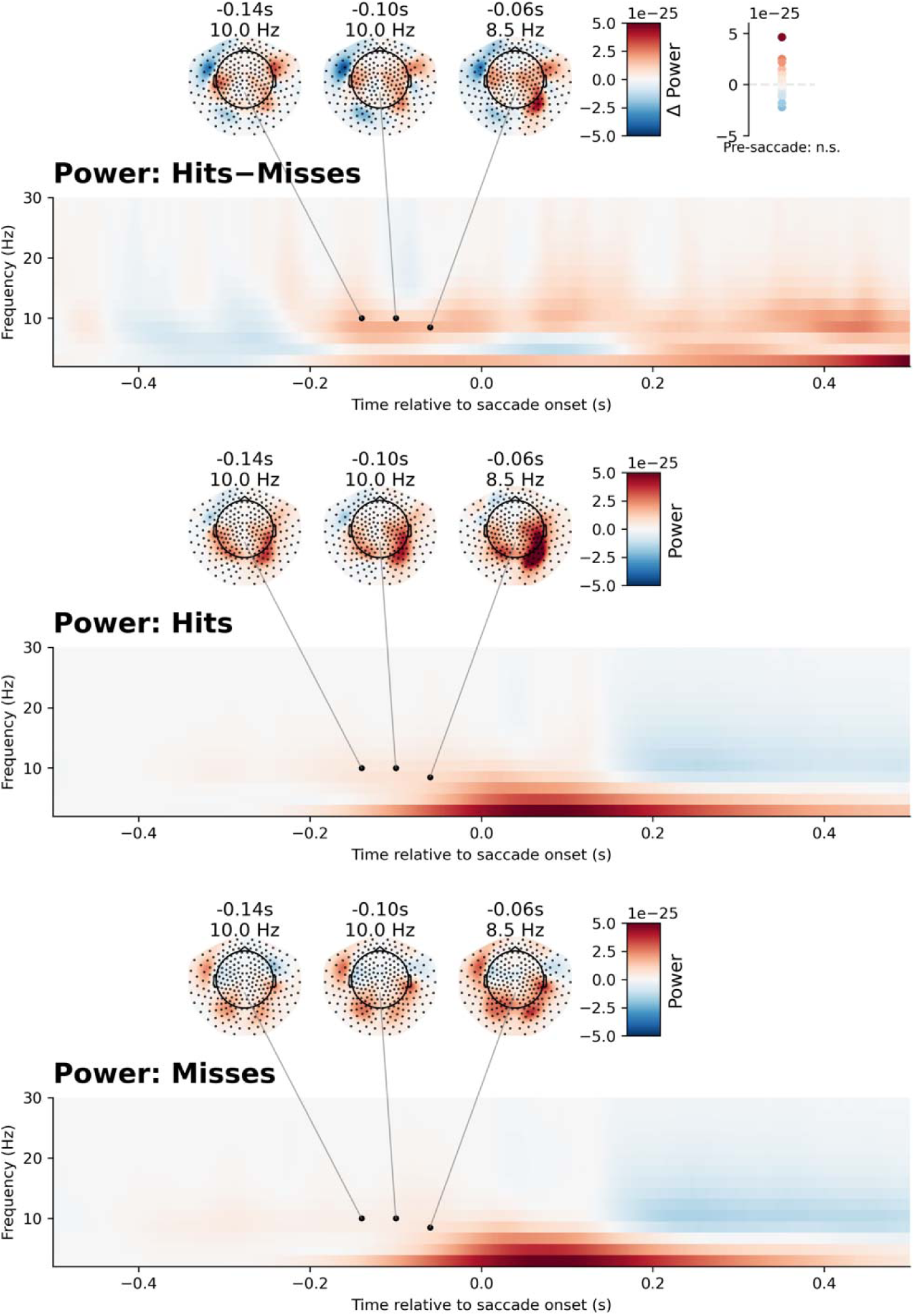
Pre-saccade differences in power, for subsequent Hits (middle), Misses (bottom), and the difference between the two (top). Although there was numerically greater alpha power preceding saccades to subsequent hits, cluster based permutation testing failed to find a statistically significant difference.

**Supplementary Figure 2.**
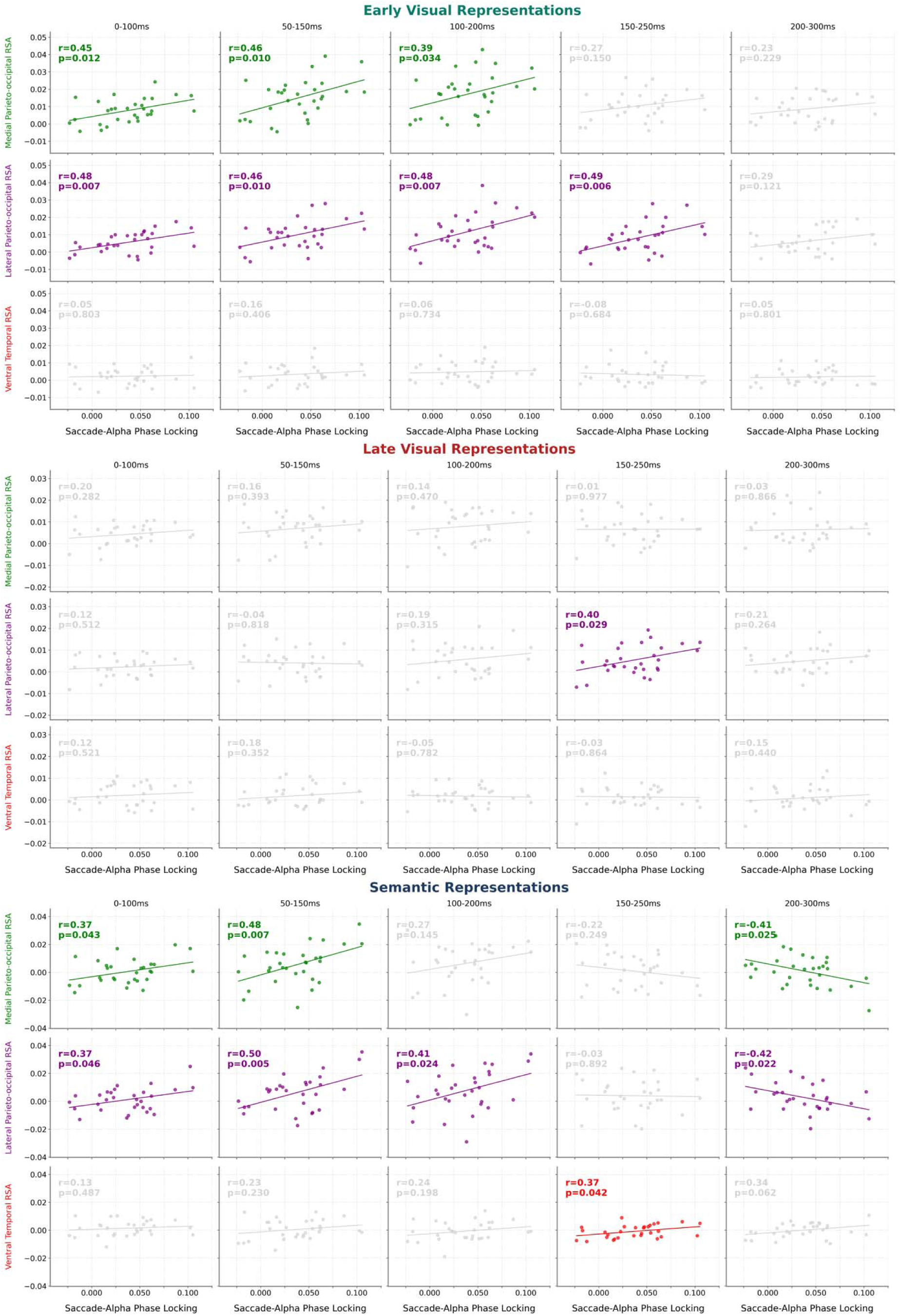
The relationship between saccade-alpha phase-locking and all three levels of post-saccade representations in left hemisphere ROIs. The top and bottom panels are the same as those shown in Figure 8. Additionally included here is the relationship with late visual representations. Colored scatter plots and lines of best fit indicate spatiotemporal windows in which there was a Pearson correlation between pre-saccade phase-locking and post-saccade representation, corresponding to a p-value < 0.05 (uncorrected).

**Supplementary Figure 3.**
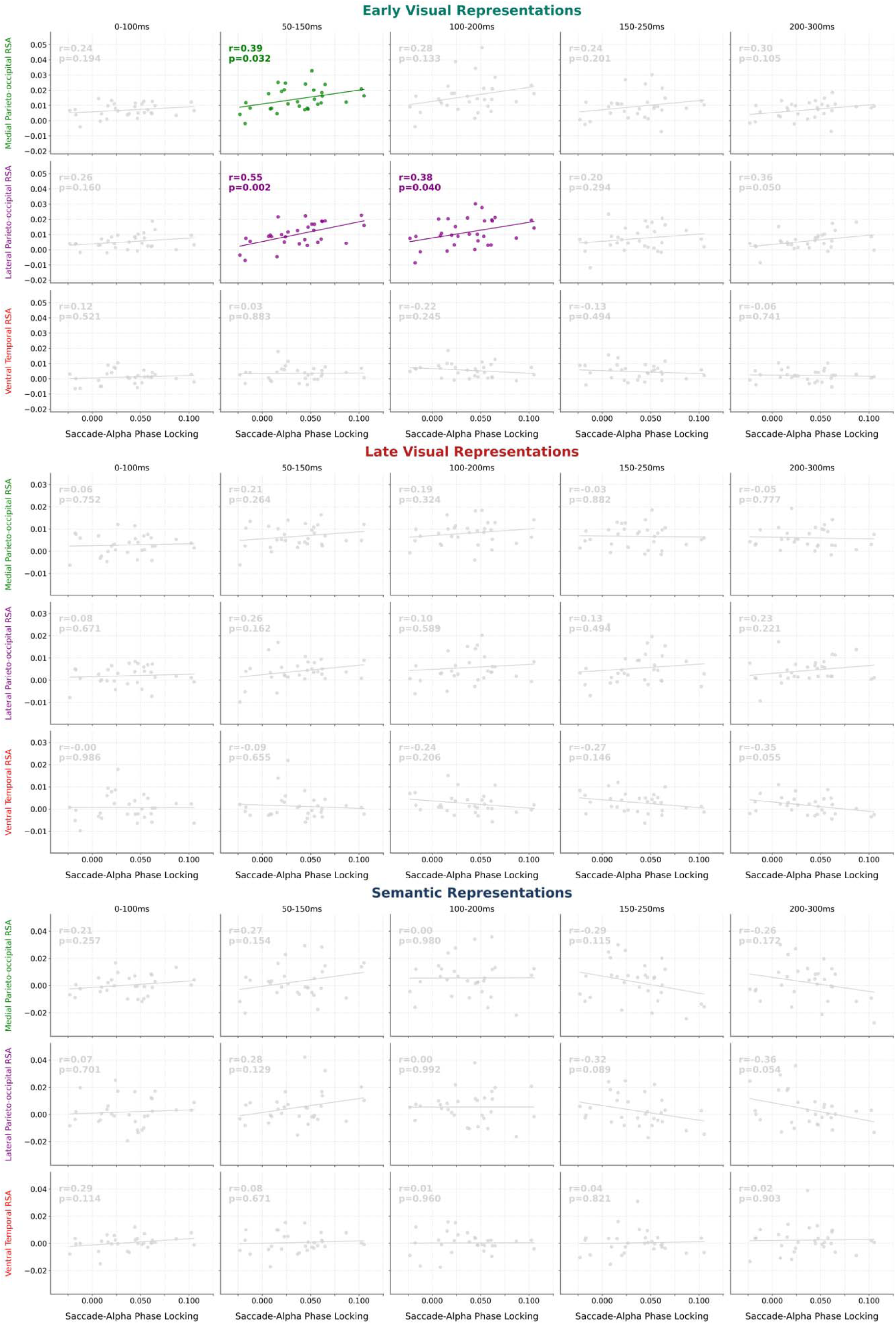
The relationship between saccade-alpha phase-locking and all three levels of post-saccade representations in right hemisphere ROIs. Colored scatter plots and lines of best fit indicate spatiotemporal windows in which there was a Pearson correlation between pre-saccade phase-locking and post-saccade representation, corresponding to a p-value < 0.05 (uncorrected).

## BIBLIOGRAPHY

Amme, C., Sulewski, P., Spaak, E., Hebart, M. N., König, P., & Kietzmann, T. C. (2024). Saccade onset, not fixation onset, best explains early responses across the human visual cortex during naturalistic vision (p. 2024.10.25.620167). bioRxiv. 10.1101/2024.10.25.620167

Auerbach-Asch, C. R., & Deouell, L. Y. (2024). Beyond Stimulus Onset: Ongoing Fixations Within an Object Do Not Re-evoke Category Representations During Free-Viewing (p. 2024.12.02.623991). bioRxiv. 10.1101/2024.12.02.623991

Berger, H. (1929). Über das elektroenkephalogramm des menschen. Archiv für psychiatrie und nervenkrankheiten, 87(1), 527–570.

Busch, N. A., Dubois, J., & VanRullen, R. (2009). The Phase of Ongoing EEG Oscillations Predicts Visual Perception. Journal of Neuroscience, 29(24), 7869–7876. 10.1523/JNEUROSCI.0113-09.2009

Carlson, T. A., Hogendoorn, H., Kanai, R., Mesik, J., & Turret, J. (2011). High temporal resolution decoding of object position and category. Journal of Vision, 11(10), 9. 10.1167/11.10.9

Carlson, T., Tovar, D. A., Alink, A., & Kriegeskorte, N. (2013). Representational dynamics of object vision: The first 1000 ms. Journal of Vision, 13(10), 1. 10.1167/13.10.1

Chan, J. P. K., Kamino, D., Binns, M. A., & Ryan, J. D. (2011). Can Changes in Eye Movement Scanning Alter the Age-Related Deficit in Recognition Memory? Frontiers in Psychology, 2. 10.3389/fpsyg.2011.00092

Clarke, A., Devereux, B. J., & Tyler, L. K. (2018). Oscillatory Dynamics of Perceptual to Conceptual Transformations in the Ventral Visual Pathway. Journal of Cognitive Neuroscience, 30(11), 1590–1605. 10.1162/jocn_a_01325

Cree, G. S., McNorgan, C., & McRae, K. (2006). Distinctive features hold a privileged status in the computation of word meaning: Implications for theories of semantic memory. *Journal of Experimental Psychology: Learning*, Memory, and Cognition, 32(4), 643–658. 10.1037/0278-7393.32.4.643

Dale, A. M., Fischl, B., & Sereno, M. I. (1999). Cortical Surface-Based Analysis: I. Segmentation and Surface Reconstruction. NeuroImage, 9(2), 179–194. 10.1006/nimg.1998.0395

Dale, A. M., & Sereno, M. I. (1993). Improved Localizadon of Cortical Activity by Combining EEG and MEG with MRI Cortical Surface Reconstruction: A Linear Approach. Journal of Cognitive Neuroscience, 5(2), 162–176. 10.1162/jocn.1993.5.2.162

Desikan, R. S., Ségonne, F., Fischl, B., Quinn, B. T., Dickerson, B. C., Blacker, D., Buckner, R. L., Dale, A. M., Maguire, R. P., Hyman, B. T., Albert, M. S., & Killiany, R. J. (2006). An automated labeling system for subdividing the human cerebral cortex on MRI scans into gyral based regions of interest. NeuroImage, 31(3), 968–980. 10.1016/j.neuroimage.2006.01.021

Devereux, B. J., Clarke, A., & Tyler, L. K. (2018). Integrated deep visual and semantic attractor neural networks predict fMRI pattern-information along the ventral object processing pathway. Scientific Reports, 8(1), 10636. 10.1038/s41598-018-28865-1

Dijk, H. van, Schoffelen, J.-M., Oostenveld, R., & Jensen, O. (2008). Prestimulus Oscillatory Activity in the Alpha Band Predicts Visual Discrimination Ability. Journal of Neuroscience, 28(8), 1816–1823. 10.1523/JNEUROSCI.1853-07.2008

Drewes, J., & VanRullen, R. (2011). This Is the Rhythm of Your Eyes: The Phase of Ongoing Electroencephalogram Oscillations Modulates Saccadic Reaction Time. The Journal of Neuroscience, 31(12), 4698–4708. 10.1523/JNEUROSCI.4795-10.2011

Dugué, L., Marque, P., & VanRullen, R. (2011). The Phase of Ongoing Oscillations Mediates the Causal Relation between Brain Excitation and Visual Perception. Journal of Neuroscience, 31(33), 11889–11893. 10.1523/JNEUROSCI.1161-11.2011

Engemann, D. A., & Gramfort, A. (2015). Automated model selection in covariance estimation and spatial whitening of MEG and EEG signals. NeuroImage, 108, 328–342. 10.1016/j.neuroimage.2014.12.040

Ergenoglu, T., Demiralp, T., Bayraktaroglu, Z., Ergen, M., Beydagi, H., & Uresin, Y. (2004). Alpha rhythm of the EEG modulates visual detection performance in humans. Cognitive Brain Research, 20(3), 376–383. 10.1016/j.cogbrainres.2004.03.009

Fakche, C., Hickey, C., & Jensen, O. (2024). Fast Feature- and Category-Related Parafoveal Previewing Support Free Visual Exploration. The Journal of Neuroscience, 44(49), e0841242024. 10.1523/JNEUROSCI.0841-24.2024

Fischl, B. (2012). FreeSurfer. NeuroImage, 62(2), 774–781. 10.1016/j.neuroimage.2012.01.021

Fischl, B., Sereno, M. I., Tootell, R. B. H., & Dale, A. M. (n.d.). High resolution intersubject averaging and a coordinate system for the cortical surface. Retrieved September 30, 2025, from https://onlinelibrary.wiley.com/doi/10.1002/(SICI)1097-0193(1999)8:4<272::AID-HBM10>3.0.CO;2-4

Fischl, B., van der Kouwe, A., Destrieux, C., Halgren, E., Ségonne, F., Salat, D. H., Busa, E., Seidman, L. J., Goldstein, J., Kennedy, D., Caviness, V., Makris, N., Rosen, B., & Dale, A. M. (2004). Automatically Parcellating the Human Cerebral Cortex. Cerebral Cortex, 14(1), 11–22. 10.1093/cercor/bhg087

Flick, G., Abdullah, O., & Pylkkänen, L. (2021). From letters to composed concepts: A magnetoencephalography study of reading. Human Brain Mapping, 42(15), 5130–5153. 10.1002/hbm.25608

Gips, B., Van Der Eerden, J. P. J. M., & Jensen, O. (2016). A biologically plausible mechanism for neuronal coding organized by the phase of alpha oscillations. European Journal of Neuroscience, 44(4), 2147–2161. 10.1111/ejn.13318

Gwilliams, L., Lewis, G. A., & Marantz, A. (2016). Functional characterisation of letter-specific responses in time, space and current polarity using magnetoencephalography. NeuroImage, 132, 320–333. 10.1016/j.neuroimage.2016.02.057

Hanslmayr, S., Klimesch, W., Sauseng, P., Gruber, W., Doppelmayr, M., Freunberger, R., & Pecherstorfer, T. (2005). Visual discrimination performance is related to decreased alpha amplitude but increased phase locking. Neuroscience Letters, 375(1), 64–68. 10.1016/j.neulet.2004.10.092

Hanslmayr, S., Volberg, G., Wimber, M., Dalal, S. S., & Greenlee, M. W. (2013). Prestimulus Oscillatory Phase at 7 Hz Gates Cortical Information Flow and Visual Perception. Current Biology, 23(22), 2273–2278. 10.1016/j.cub.2013.09.020

Hovhannisyan, M., Clarke, A., Geib, B. R., Cicchinelli, R., Monge, Z., Worth, T., Szymanski, A., Cabeza, R., & Davis, S. W. (2021). The visual and semantic features that predict object memory: Concept property norms for 1,000 object images. Memory & Cognition, 49(4), 712–731. 10.3758/s13421-020-01130-5

Jensen, O., Gips, B., Bergmann, T. O., & Bonnefond, M. (2014). Temporal coding organized by coupled alpha and gamma oscillations prioritize visual processing. Trends in Neurosciences, 37(7), 357–369. 10.1016/j.tins.2014.04.001

Jensen, O., & Mazaheri, A. (2010). Shaping Functional Architecture by Oscillatory Alpha Activity: Gating by Inhibition. Frontiers in Human Neuroscience, 4. 10.3389/fnhum.2010.00186

Jensen, O., Pan, Y., Frisson, S., & Wang, L. (2021). An oscillatory pipelining mechanism supporting previewing during visual exploration and reading. Trends in Cognitive Sciences, 25(12), 1033–1044. 10.1016/j.tics.2021.08.008

Kragel, J. E., Schuele, S., VanHaerents, S., Rosenow, J. M., & Voss, J. L. (2021). Rapid coordination of effective learning by the human hippocampus. Science Advances, 7(25), eabf7144. 10.1126/sciadv.abf7144

Kriegeskorte, N., Mur, M., & Bandettini, P. A. (2008). Representational similarity analysis— Connecting the branches of systems neuroscience. Frontiers in Systems Neuroscience, 2. 10.3389/neuro.06.004.2008

Krizhevsky, A., Sutskever, I., & Hinton, G. E. (2017). ImageNet classification with deep convolutional neural networks. Commun. ACM, 60(6), 84–90. 10.1145/3065386

Ledoit, O., & Wolf, M. (2004). A well-conditioned estimator for large-dimensional covariance matrices. Journal of Multivariate Analysis, 88(2), 365–411. 10.1016/S0047-259X(03)00096-4

Lin, F.-H., Belliveau, J. W., Dale, A. M., & Hämäläinen, M. S. (2006). Distributed current estimates using cortical orientation constraints. Human Brain Mapping, 27(1), 1–13. 10.1002/hbm.20155

Liu, B., Nobre, A., & van Ede, F. (2022). Relating microsaccades and EEG-alpha activity during covert spatial attention in visual working memory. Journal of Vision, 22(14), 3472. 10.1167/jov.22.14.3472

Loftus, G. R. (1972). Eye fixations and recognition memory for pictures. Cognitive Psychology, 3(4), 525–551. 10.1016/0010-0285(72)90021-7

Maris, E., & Oostenveld, R. (2007). Nonparametric statistical testing of EEG- and MEG-data. Journal of Neuroscience Methods, 164(1), 177–190. 10.1016/j.jneumeth.2007.03.024

Mathewson, K. E., Gratton, G., Fabiani, M., Beck, D. M., & Ro, T. (2009). To See or Not to See: Prestimulus α Phase Predicts Visual Awareness. Journal of Neuroscience, 29(9), 2725– 2732. 10.1523/JNEUROSCI.3963-08.2009

Mathewson, K. E., Lleras, A., Beck, D. M., Fabiani, M., Ro, T., & Gratton, G. (2011). Pulsed Out of Awareness: EEG Alpha Oscillations Represent a Pulsed-Inhibition of Ongoing Cortical Processing. Frontiers in Psychology, 2. 10.3389/fpsyg.2011.00099

Muttenthaler, L., & Hebart, M. N. (2021). THINGSvision: A Python Toolbox for Streamlining the Extraction of Activations From Deep Neural Networks. Frontiers in Neuroinformatics, 15. 10.3389/fninf.2021.679838

Nikolaev, A. R., Jurica, P., Nakatani, C., Plomp, G., & van Leeuwen, C. (2013). Visual encoding and fixation target selection in free viewing: Presaccadic brain potentials. Frontiers in Systems Neuroscience, 7. 10.3389/fnsys.2013.00026

Nikolaev, A. R., Meghanathan, R. N., & van Leeuwen, C. (2016). Combining EEG and eye movement recording in free viewing: Pitfalls and possibilities. Brain and Cognition, 107, 55–83. 10.1016/j.bandc.2016.06.004

Olsen, R. K., Moses, S. N., Riggs, L., & Ryan, J. D. (2012). The hippocampus supports multiple cognitive processes through relational binding and comparison. Frontiers in Human Neuroscience, 6. 10.3389/fnhum.2012.00146

Olsen, R. K., Rondina II, R., Riggs, L., Meltzer, J. A., & Ryan, J. D. (2013). Hippocampal and neocortical oscillatory contributions to visuospatial binding and comparison. Journal of Experimental Psychology: General, 142(4), 1335–1345. 10.1037/a0034043

Olsen, R. K., Sebanayagam, V., Lee, Y., Moscovitch, M., Grady, C. L., Rosenbaum, R. S., & Ryan, J. D. (2016). The relationship between eye movements and subsequent recognition: Evidence from individual differences and amnesia. Cortex, 85, 182–193. 10.1016/j.cortex.2016.10.007

Pan, Y., Popov, T., Frisson, S., & Jensen, O. (2023). Saccades are locked to the phase of alpha oscillations during natural reading. PLOS Biology, 21(1), e3001968. 10.1371/journal.pbio.3001968

Peirce, J., Gray, J. R., Simpson, S., MacAskill, M., Höchenberger, R., Sogo, H., Kastman, E., & Lindeløv, J. K. (2019). PsychoPy2: Experiments in behavior made easy. Behavior Research Methods, 51(1), 195–203. 10.3758/s13428-018-01193-y

Popov, T., Miller, G. A., Rockstroh, B., Jensen, O., & Langer, N. (2021). Alpha oscillations link action to cognition: An oculomotor account of the brain’s dominant rhythm (p. 2021.09.24.461634). bioRxiv. 10.1101/2021.09.24.461634

Popov, T., & Staudigl, T. (2023). Cortico-ocular coupling in the service of episodic memory formation. Progress in Neurobiology, 227, 102476. 10.1016/j.pneurobio.2023.102476

Ryan, J. D., Shen, K., & Liu, Z.-X. (2020). The intersection between the oculomotor and hippocampal memory systems: Empirical developments and clinical implications. Annals of the New York Academy of Sciences, 1464(1), 115–141. 10.1111/nyas.14256

Staudigl, T., Hartl, E., Noachtar, S., Doeller, C. F., & Jensen, O. (2017). Saccades are phase-locked to alpha oscillations in the occipital and medial temporal lobe during successful memory encoding. PLOS Biology, 15(12), e2003404. 10.1371/journal.pbio.2003404

Tallon-Baudry, C., Bertrand, O., Delpuech, C., & Pernier, J. (1996). Stimulus Specificity of Phase-Locked and Non-Phase-Locked 40 Hz Visual Responses in Human. Journal of Neuroscience, 16(13), 4240–4249. 10.1523/JNEUROSCI.16-13-04240.1996

Thickbroom, G. W., & Mastaglia, F. L. (1985). Presaccadic ‘spike’ potential: Investigation of topography and source. Brain Research, 339(2), 271–280. 10.1016/0006-8993(85)90092-7

Tuladhar, A. M., Huurne, N. ter, Schoffelen, J.-M., Maris, E., Oostenveld, R., & Jensen, O. (2007). Parieto-occipital sources account for the increase in alpha activity with working memory load. Human Brain Mapping, 28(8), 785–792. 10.1002/hbm.20306

van Diepen, R. M., & Mazaheri, A. (2018). The Caveats of observing Inter-Trial Phase-Coherence in Cognitive Neuroscience. Scientific Reports, 8(1), 2990. 10.1038/s41598-018-20423-z

Voloh, B., & Womelsdorf, T. (2016). A Role of Phase-Resetting in Coordinating Large Scale Neural Networks During Attention and Goal-Directed Behavior. Frontiers in Systems Neuroscience, 10. 10.3389/fnsys.2016.00018

von Seth, J., Nicholls, V. I., Tyler, L. K., & Clarke, A. (2023). Recurrent connectivity supports higher-level visual and semantic object representations in the brain. Communications Biology, 6(1), 1207. 10.1038/s42003-023-05565-9

Voss, J. L., Bridge, D. J., Cohen, N. J., & Walker, J. A. (2017). A Closer Look at the Hippocampus and Memory. Trends in Cognitive Sciences, 21(8), 577–588. 10.1016/j.tics.2017.05.008

Wu, X., Popov, T., Beilner, T., Fearns, N., Vollmar, C., Kaufmann, E., Remi, J., & Staudigl, T. (2025). Low-frequency brain oscillations reflect the dynamics of the oculomotor system: A new perspective on subsequent memory effects (p. 2025.07.29.667451). bioRxiv. 10.1101/2025.07.29.667451

